# TDP-43 toxic gain of function links ALS, FTD and Alzheimer’s disease via mis-splicing

**DOI:** 10.1101/2025.04.20.648873

**Authors:** Welmoed van Zuiden, Thea Meimoun, Chen Bar, Aviad Siany, Bar Abarbanel, Lihi Moshe, Nancy S Yacovzada, Emmanuel Amzallag, Aleksandar Rajkovic, Liat Alyagor, Eviatar Weizman, Manuela Neumann, Colin Smith, Karina McDade, Andrew Strange, Rodrigo Siqueira Kazu, Marcel Weinreich, Aron S Buchman, Yanling Wang, David A Bennett, Bingqing Zhao, Lixia Jiang, Joanna Bi, Michael P Snyder, Dmytro Morderer, Jorge Garcia Morato, Wilfried Rossoll, Madison M Reeves, Dennis W. Dickson, Mercedes Prudencio, Peter T Nelson, Jonathan D Glass, Adam N Trautwig, Mingee Chung, Zachary T McEachin, Nicholas T Seyfried, Johnathan Cooper-Knock, Eran Hornstein

## Abstract

TDP-43 proteinopathy is a hallmark of amyotrophic lateral sclerosis (ALS) and frontotemporal lobar degeneration (FTLD) ^1^. It is also common in Alzheimer’s disease (AD) and limbic-predominant age-related TDP-43 encephalopathy (LATE) ^2,3^, but its functional consequences are unclear. Here, we show that TDP-43 pathology in AD is broadly associated with splicing abnormalities. We define a conserved TDP-43-dependent splicing signature in AD that includes AD-relevant mis-spliced mRNAs such as amyloid precursor protein (*APP*). TDP-43 drives the formation of elongated *APP* isoforms across ALS, FTLD-TDP and AD, providing a mechanism for the long-standing observation of *APP* isoform dysregulation ^4^. *APP* isoform imbalance correlates with and partially mediates the association between TDP-43 pathology and Aβ burden and is spatially associated with pTDP-43 and mis-spliced *STMN2* in LATE hippocampus. Contrary to a pure nuclear loss-of-function model ^5,6^, mis-splicing is driven by cytoplasmic TDP-43 gain-of-function that co-sequesters splicing-regulatory RNA-binding proteins. Knockdown of TDP-43-associated RNA-binding proteins recapitulates disease-relevant splicing changes in AD and ALS/FTLD-TDP, and some of these proteins colocalize with cytoplasmic pTDP-43 in neuropathological samples. Our findings establish cytoplasmic TDP-43-driven splicing dysfunction as a key component of neurodegeneration, define disease-relevant mis-spliced mRNAs in AD and unite AD with ALS/FTLD-TDP through a shared mechanism of splicing factor sequestration.

## Introduction

Splicing dysregulation is implicated in several neurodegenerative disorders, including ALS, FTLD and AD ^7^. TDP-43, encoded by the *TARDBP* gene, is a ubiquitous nuclear DNA- and RNA-binding protein (RBP) that regulates RNA processing, including splicing ^8^. In nearly all cases of ALS and up to 50% of FTLD patients, TDP-43 forms insoluble cytoplasmic inclusions and is depleted from the nucleus ^1^, leading to widespread splicing disruption, including cryptic exon inclusion in mRNAs such as cytoskeleton-related *STMN2* or synaptic *UNC13A* ^5,6,8–12^.

TDP-43 proteinopathy is also found in 50-75% of AD neuropathology specimens ^2^ and is the defining feature of a common comorbidity in the aging brain. In AD, TDP-43 proteinopathy stereotypically spreads from the amygdala and hippocampus to the neocortex, correlates with cognitive decline and pathology severity, and is associated with increased Tau burden ^13,14^. While aberrant splicing is a recognized feature of AD ^15^, the contribution of TDP-43 to these defects remains largely unexplored ^16–18^.

The proteolytic processing of amyloid precursor protein (APP) into the neurotoxic Aβ peptide is thought to be a crucial step in Alzheimer’s disease (AD) ^19^. Alternative splicing of exons 7 and 8 produces three major *APP* isoforms conserved across species: the neuron-enriched APP695 and the longer APP751 and APP770. APP751 and APP770 contain the Kunitz protease inhibitor (KPI) domain, and APP770 also contains the OX-2 domain. KPI(+)APP protein and mRNA levels are elevated in AD and associated with increased Aβ production ^4^. However, the mechanism of *APP* alternative splicing is poorly understood.

Here, we demonstrate that TDP-43 pathology is broadly associated with splicing abnormalities in AD, including *APP* alternative splicing, suggesting a mechanistic commonality with ALS/FTLD-TDP. We reveal a conserved TDP-43-dependent splicing signature across brain regions and AD cohorts, provide a mechanism for *APP* isoform dysregulation first reported decades ago ^4^, and show that it partially mediates Aβ pathology. Surprisingly, a TDP-43 toxic gain-of-function, rather than nuclear loss-of-function, drives these shifts in *APP* and other mRNA isoforms, involving cytoplasmic co-sequestration of additional RBPs that colocalize with pTDP-43 in neuropathological samples. Collectively, TDP-43 pathology links seemingly disparate neurodegenerative diseases via shared splicing dysfunction.

## Results

### TDP-43 splicing abnormalities in Alzheimer’s disease

To investigate whether TDP-43 pathology contributes to dysregulated splicing in AD, we analyzed RNA-seq data of postmortem dorsolateral prefrontal cortex (DLPFC) from the Religious Orders Study and Rush Memory and Aging Project (ROSMAP ^20^, Synapse: syn23446022). Using MAJIQ ^21^, we compared patients clinically diagnosed with AD dementia (*n =* 236) to controls without cognitive impairment (*n =* 200), and AD brains with TDP-43 pathology (*n =* 110) to those without (*n =* 126) (**Supplementary Table 1**). Aberrant splicing was found in 2354 mRNAs in AD versus controls, and in 963 mRNAs in AD with versus without TDP-43 pathology (|ΔPSI| > 0.02, signifying the change in percent spliced in (PSI) between conditions, and *P* < 0.05; **Supplementary Table 2**). A significant portion of mRNAs aberrantly spliced in AD was also mis-spliced in AD with TDP-43 pathology (Fisher’s exact, *P* < 0.0001, **Figure 1A**), including most mRNAs previously reported as mis-spliced in AD ^15^. Comparison of TDP-43-positive and TDP-43-negative AD samples versus controls suggests that ∼20% of AD-associated splicing changes are TDP-43-dependent (**Extended Data Figure 1**).

**Figure 1:**
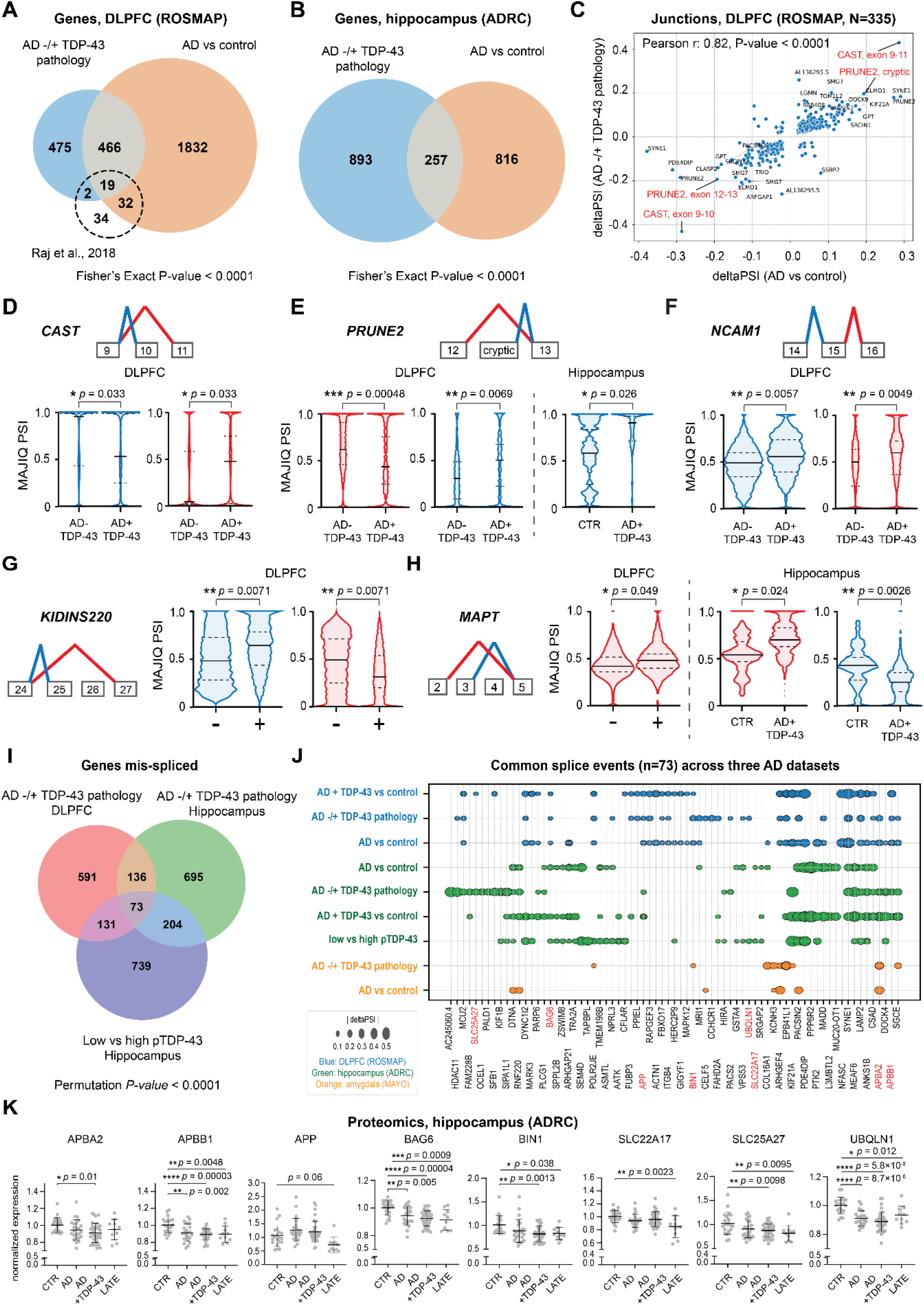
TDP-43 pathology is associated with widespread mis-splicing in Alzheimer’s disease. Overlap of TDP-43-associated mis-splicing in AD bulk RNA-seq (MAJIQ; |ΔPSI| > 0.02, P < 0.05): **(A)** ROSMAP DLPFC: AD versus cognitively unimpaired controls (*n =* 236, 200; orange) and AD with versus without TDP-43 pathology (AD+TDP-43, *n =* 110; AD-TDP-43, *n =* 126; blue); **(B)** ADRC hippocampus^23^: AD versus controls (n = 60, 21; orange) and AD+TDP-43 versus AD-TDP-43 (*n =* 35, *n =* 25; blue). One-sided Fisher’s exact test, *P* < 0.0001. Only mRNAs detected in both analyses were included (1073 of 1080 for AD versus controls; all 1150 for AD with versus without TDP-43 pathology). **(C)** Junction-level ΔPSI in ROSMAP, as in (A): AD versus controls (x-axis) and AD+TDP-43 versus AD-TDP-43 (y-axis). mRNAs with |ΔPSI| > 0.15 in either comparison are labeled. Two-sided Pearson’s correlation *r* = 0.82, *P* < 0.0001. **(D-H)** Splice junction diagrams of PSI for *CAST* exons 9-11 **(D)**, *PRUNE2* exons 12-13 **(E)**, *NCAM1* exons 14-16 **(F)**, *KIDINS220* exons 24-27 **(G)** and *MAPT* exons 2-5 **(H)** in ROSMAP DLPFC (AD+TDP-43 versus AD-TDP-43; all cases with versus without TDP-43 pathology (+, −)) and in ADRC hippocampus (AD+TDP-43 versus control). Violin plots show median and interquartile range; colors correspond to the junctions in the diagram and only cases with detectable junction PSI are included. Two-sided TNOM test (MAJIQ Heterogen), not adjusted for multiple comparisons. **(I)** High-confidence mis-spliced mRNAs shared between DLPFC (ROSMAP) and hippocampal (ADRC) AD+TDP-43 versus AD-TDP-43 (*n* as in A,B), and ADRC samples stratified by quantitative pTDP-43 immunoassay ^23^ (lowest 25% versus highest 75%; *n =* 25, 66). One-sided permutation test (10,000 permutations), *P* < 1 × 10⁻⁴ (minimum attainable). **(J)** Bubble heat map of |ΔPSI| values for the 73 high-confidence mRNAs in (I) ROSMAP DLPFC, ADRC hippocampus, and Mayo Clinic amygdala (*n =* 68 AD (49 TDP-43+, 19 TDP-43-), *n =* 17 controls). Junctions matched by unique MAJIQ local splicing variation identifiers. Red: genes associated with AD. **(K)** Hippocampal protein levels (mass spectrometry ^71,72^, normalized to control) of the eight genes in red in (J); control, AD, AD+TDP-43, LATE (*n* = 21, 24, 35, 10 individuals respectively). Data are presented as mean ± s.d. One-way ANOVA with two-sided Dunnett’s multiple-comparisons test. (D-H,K) Exact P values are shown in the figure. \**P* < 0.05, \*\**P* < 0.01, \*\*\**P* < 0.001, \*\*\*\**P* < 0.0001.

As TDP-43 pathology in AD is less prominent in the frontal cortex than in limbic regions ^22^, we also analyzed hippocampal tissue from the University of Kentucky Alzheimer’s Disease Research Center (ADRC)^23–25^. These specimens were characterized by Alzheimer’s disease neuropathological changes and limbic-predominant age-related TDP-43 encephalopathy neuropathological changes (ADNC + LATE-NC). We identified 1080 genes mis-spliced in AD versus controls and 1150 in AD with versus without TDP-43 pathology (|ΔPSI| > 0.02, *P* < 0.05; **Supplementary Table 2**), with significant overlap (Fisher’s exact test, *P* < 0.0001; **Figure 1B**).

In the ROSMAP DLPFC, 1623 exact splice junctions were associated with TDP-43 pathology and 6413 with AD, 335 of them shared (Fisher’s exact test, *P* < 0.0001). The magnitude and direction of splicing changes at these shared junctions were highly concordant (Pearson *r* = 0.82, *P* < 0.0001, **Figure 1C**).

Among the most misregulated RNAs in AD with versus without TDP-43 pathology were the proposed AD susceptibility genes *CAST* (exon 10 inclusion decreased by 43%) and *PRUNE2* (inclusion of cryptic exon between exons 12 and 13 increased by 20%-35%; **Figure 1D,E**). Other highly mis-spliced TDP-43-dependent targets included *NCAM1* and *KIDINS220* (**Figure 1F,G**), both implicated in AD.

Alternative splicing of microtubule-associated protein Tau (*MAPT*) mRNA is predicted to yield over 30 isoforms, primarily through inclusion of exons 2, 3, and 10 ^26^. The exon 2-5 junction was used 5% more in ROSMAP DLPFC cases with TDP-43 pathology than in controls (*P* < 0.05, **Figure 1H**). Similarly, in hippocampal AD specimens with TDP-43 pathology (LATE-NC + ADNC), exon 2-5 junction use increased by 16% (*P* < 0.05), while exon 3-5 junction use decreased by 18% (*P* < 0.01), suggesting that TDP-43 pathology contributes to aberrant *MAPT* splicing in the human brain.

Next, we defined a high-confidence set of TDP-43-associated mRNA targets by intersecting mRNAs mis-spliced in DLPFC and hippocampal AD samples with versus without TDP-43 pathology and in hippocampal AD samples stratified by pTDP-43 burden. This revealed 73 genes consistently mis-spliced across all three datasets, out of 2569 spliced genes identified in all conditions (**Figure 1I**). This overlap far exceeds chance: in a permutation test in which gene sets of equal size were randomly sampled 10,000 times from the common background, the three-way overlap was significantly enriched (*P* < 0.0001). These TDP-43-associated splicing alterations are therefore conserved across brain regions and analytical contrasts.

To assess cross-dataset consistency, we examined splice junction-level values for these 73 mRNAs across DLPFC (ROSMAP), hippocampus (ADRC), and an independent amygdala RNA-seq dataset (Mayo Clinic). Most shared splice events showed concordant magnitude across all three datasets (|ΔPSI|, **Figure 1J**), despite differences in tissue composition, sequencing depth, and cohort structure. These results define a conserved TDP-43-dependent splicing program in AD, while reflecting the heterogeneity of splicing regulation across brain regions.

Several mis-spliced mRNAs within this signature encode proteins with suggested genetic or functional links to AD, including (ΔPSI in hippocampus, TDP-43-positive AD vs control, unless stated otherwise) *APBA2* (exon 8 to 9, +29%), *APBB1* (exon 8 to 9, +14%), *APP* (exon 6 to 9, −11%), *BAG6* (exon 23 to 25, +10%), *BIN1* (exon 12 to 13, −11%, TDP-43-positive vs TDP-43-negative AD, DLPFC), *SLC22A17* (intron between exon 6 and 7, +5%, high vs low pTDP-43), *SLC25A27* (intron between exon 2 and 3, +37%, TDP-43-positive vs TDP-43-negative AD) and *UBQLN1* (exon 8 to 9, −7%, high vs low pTDP-43), implying potential functional consequences for disease-relevant pathways.

Finally, to test whether mis-splicing alters protein abundance, we examined hippocampal proteomics for these eight AD-relevant high-confidence targets, which displayed significant changes in abundance across disease or TDP-43 pathology status (**Figure 1K**). Together, these results demonstrate that TDP-43 pathology is associated with widespread, quantitative, and reproducible splicing abnormalities in AD, supporting TDP-43 dysfunction as a major driver of transcriptomic and proteomic dysregulation.

### TDP-43 pathology drives *APP* mis-splicing

Dysregulated APP splicing has been recognized in AD for more than three decades ^4,27^, but its mechanism remains unknown. We observed significant *APP* mis-splicing in AD brains and in cases with confirmed TDP-43 pathology (**Figure 2A-D**). Specifically, the exon 6-9 junction, which produces neuronal APP695 (**Figure 2A**), was used 5-8% less in AD and TDP-43-positive samples than in controls. This was accompanied by a reciprocal 4-8% increase in inclusion of exon 7 or exons 7 and 8, producing the AD-associated APP751 and APP770 isoforms, respectively (**Figure 2B-D; Supplementary Table 3**). Notably, no *APP* mis-splicing was detected in AD samples lacking TDP-43 pathology (**Extended Data Figure 2A**), identifying TDP-43 as the essential driver of these changes. In hippocampal samples, *APP* mis-splicing likewise correlated strictly with pTDP-43 pathology and higher pTDP-43 concentrations (**Figure 2E, F; Extended Data Figure 2C, D**).

**Figure 2:**
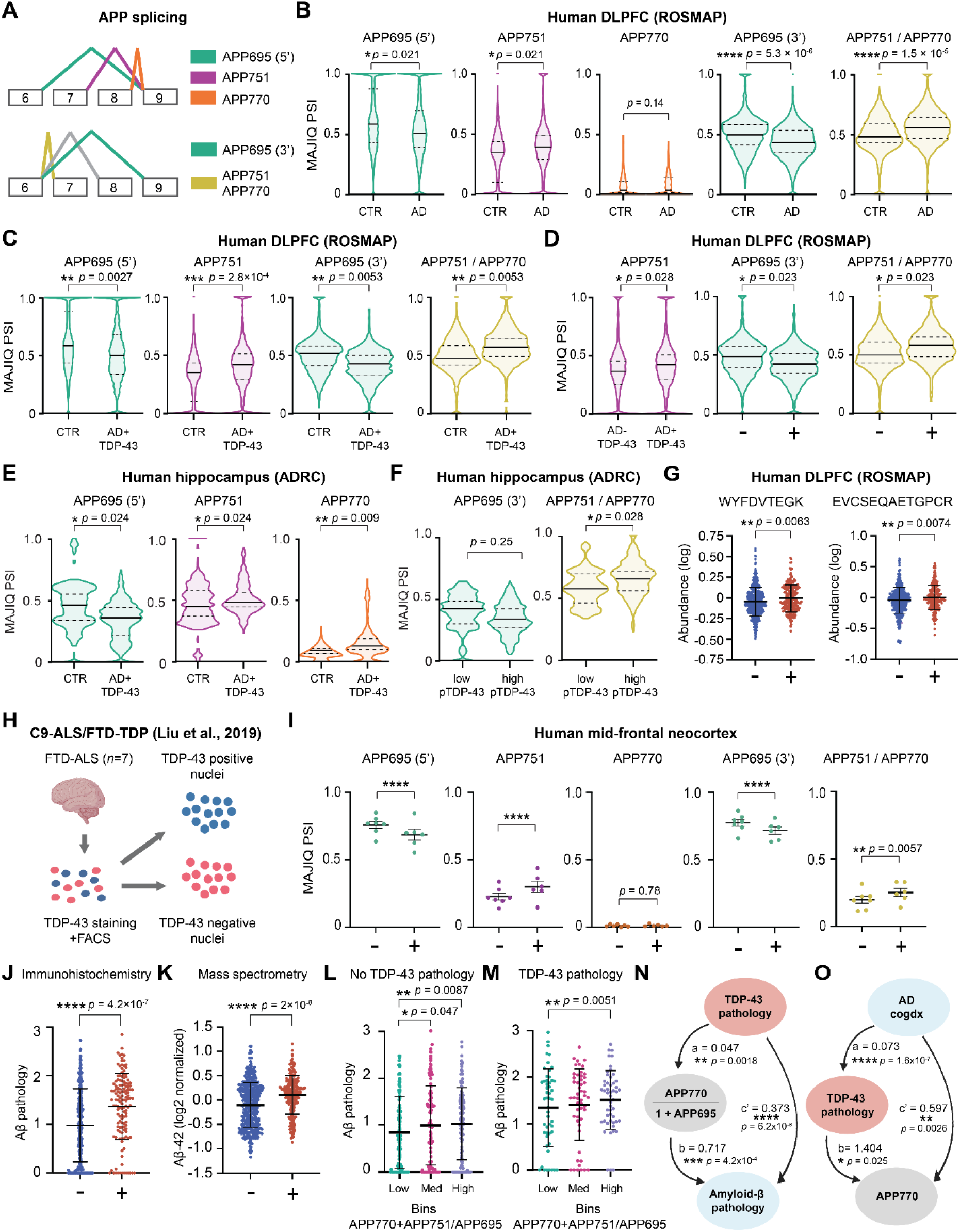
TDP-43 pathology and consequent *APP* mis-splicing lead to increased Aβ production. **(A)** *APP* exons 6-9 splice junctions and corresponding isoforms. *APP* PSI in ROSMAP DLPFC: **(B)** AD (*n* = 236) versus controls (CTR, *n* = 200), **(C)** AD+TDP-43 (*n* = 110) versus controls and **(D)** AD+TDP-43 versus AD-TDP-43 (*n* = 126), and all samples with TDP-43 pathology (*n* = 153) versus without (*n* = 349) TDP-43 pathology. *APP* PSI in ADRC hippocampus: **(E)** AD+TDP-43 (*n* = 35) versus controls (*n* = 21) or **(F)** samples with the lowest 25% (*n =* 25) versus the highest 75% (*n =* 66) of pTDP-43 immunoassay values. **B–F**, groups were compared with the two-sided TNOM test (MAJIQ Heterogen); median PSI and interquartile range are shown, and only cases with detectable junction-specific PSI are included. **(G)** Abundance of APP751-and APP770-specific peptides (WYFDVTEGK, EVCSEQAETGPCR) with (+; n = 216, 217) or without (-; n = 424, 427) TDP-43 pathology (data from Seifar et al ^28^, TAMPOR-normalized, log2). Unpaired two-sided t-test after ROUT outlier removal. **(H)** Diagram of *C9orf72* ALS/FTLD TDP-43-positive (no pathology, −) and TDP-43-negative (pathological, +) nuclei fractions sorting by Liu et al. ^29^ and **(I)** corresponding *APP* exons 6-9 splice junctions PSI (*n* = 7 cases, 35,000-100,000 nuclei pre-sorting per case). Mean ± s.e.m. MAJIQ ΔPSI (P = 1 − posterior probability of |ΔΨ| > 0.02); **** indicates a MAJIQ *P* < 1 × 10⁻⁴, the smallest value MAJIQ reports. Aβ pathology in ROSMAP DLPFC samples, comparing samples with or without TDP-43 as measured by **(J)** immunohistochemistry ^73^ (*n* = 153, 349) or **(K)** mass spectrometry ^28^ (*n* = 221, 431, processed as in G). Unpaired two-sided t-test. Aβ pathology burden (immunohistochemistry) across low, medium, or high ratios of 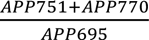 in samples **(L)** without (*n* = 349) and **(M)** with (*n* = 153) TDP-43 pathology. Two-sided Welch t-test versus the low group. Mediation analyses in ROSMAP DLPFC (*n* = 502), of **(N)** the association between TDP-43 pathology and Aβ burden partially mediated by *APP* mis-splicing 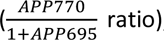, and **(O)** the association between clinical AD diagnosis (*cogdx*) and APP770 levels partially mediated by TDP-43 pathology. Unstandardized path coefficients were estimated by ordinary least squares regression without covariates (two-sided t-tests); indirect effects (a×b) were assessed by percentile bootstrap (1,000 resamples; **N**, a×b = 0.033, 95% CI 0.010–0.066; **O**, a×b = 0.102, 95% CI 0.017–0.204). *P < 0.05, **P < 0.01, ***P < 0.001, ****P < 0.0001. **G**, **J-M**, mean ± s.d. No adjustment for multiple comparisons. Exact P values are shown in the figure.

Orthogonal analysis with RNA-Seq by expectation maximization (RSEM; *n* = 502) revealed 20% lower APP695 expression (Welch’s t-test, *P* < 0.001) and higher APP751+APP770 expression (*P* < 0.01) in samples with versus without TDP-43 pathology (**Extended Data Figure 2B**). In addition, two peptides specific to the longer *APP* isoforms (APP751 and APP770) ^28^ were more abundant in ROSMAP DLPFC samples with versus without TDP-43 pathology (unpaired t-test, *P* < 0.01, **Figure 2G**) and in AD versus controls (*P* < 0.0001, **Extended Data Figure 2E**), confirming *APP* isoform dysregulation at the mRNA and protein levels.

To test whether TDP-43 pathology is associated with *APP* mis-splicing in other neurodegenerative conditions, we analyzed RNA-seq data from neuronal nuclei with TDP-43 pathology in ALS/FTLD-TDP brains from *C9orf72* repeat expansion carriers ^29^ (**Figure 2H**). Comparing TDP-43-positive to TDP-43-depleted nuclei revealed 8837 aberrant splicing events (|ΔPSI| > 0.1 and *P* < 0.1) in 2947 genes, including *UNC13A* and *STMN2* (**Supplementary Table 2**). Almost half of the TDP-43-associated mis-spliced mRNAs in AD DLPFC were also mis-spliced in ALS/FTLD-TDP (Fisher’s exact *P* < 0.0001, **Extended Data Figure 2F**), suggesting broadly shared TDP-43–dependent splicing defects across diseases. In TDP-43-depleted nuclei, usage of the APP exon 6-9 junction decreased by 10% relative to control nuclei (*P* < 0.0001, **Figure 2I** and **Supplementary Table 3**), whereas the exon 7-9 and exon 6-7 junctions were used 10% (*P* < 0.0001) and 7% (*P* < 0.01) more, respectively, resulting in increased APP751 and APP770 (**Figure 2I**). These results, from direct comparison of enriched TDP-43-positive and TDP-43-negative neuronal nuclei, extend the finding of *APP* mis-splicing downstream of TDP-43 to *C9orf72* ALS/FTLD-TDP, suggesting a shared mechanism across neurodegenerative diseases.

### Aβ is associated with TDP-43-dependent APP mis-splicing

Because elevated KPI-containing *APP* isoforms (APP751 and APP770) have been reported to correlate with increased amyloid-β (Aβ) production in AD ^30^, we examined how TDP-43 pathology and *APP* isoforms relate to Aβ in postmortem brain tissue. ROSMAP DLPFC samples with TDP-43 pathology showed ∼1.4-fold higher Aβ pathology by both immunohistochemistry (*P* < 0.0001, **Figure 2J**) and mass spectrometry ^28^ (*P*<0.001, **Figure 2K**). This remained significant when restricted to AD samples (*P* < 0.05, **Extended Data Figure 2G, H**). Moreover, binning samples by the ratio of 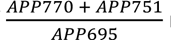 mRNA levels revealed that higher ratios significantly correlated with elevated Aβ burden (*P* < 0.05, **Figure 2L, M**). Aβ burden similarly correlated with levels of the canonical TDP-43 splicing targets, *STMN2* and *UNC13A* isoforms **(Extended Data Figure 2I-K)**, which was not identified in a previous study ^16^. These findings suggest that TDP-43 dysfunction could be a central upstream driver of Aβ pathology.

### Mediation analysis of TDP-43, *APP* and Aβ

To determine if *APP* isoform dysregulation mediates the link between TDP-43 and Aβ pathology, we performed causal mediation analysis in ROSMAP DLPFC samples (**Figure 2N**; **Supplementary Table 4**). TDP-43 pathology robustly predicted Aβ burden (total effect *c* = 0.407, *p =* 4.3 × 10⁻⁹). TDP-43 also correlated positively with inclusion of exons 7 and 8 (APP770 and APP751; *P* ≤ 0.003) and negatively with APP695 (*p* = 0.005) and strongly predicted the ratio of elongated-to-neuronal isoforms, defined as 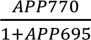. We then quantified the effect of *APP* isoforms on Aβ pathology while controlling TDP-43 (path b). 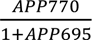 significantly attenuated the direct effect (c’) of TDP-43, mediating the largest proportion of the relationship (*ab* = 0.033, 95% CI 0.010–0.066). Individual KPI-containing isoforms also showed mediation, but APP695 alone showed no significant effect. Thus, although subtle, the 5-10% shift in *APP* splicing alters steady-state levels toward pro-amyloidogenic isoforms, suggesting one avenue by which TDP-43 contributes to Aβ accumulation.

To determine if TDP-43 pathology explains the association between clinical AD and *APP* dysregulation, we performed mediation analysis on AD diagnosis (*cogdx*), pTDP-43, and APP770 levels (**Figure 2O; Supplementary Table 4**). Consistent with previous reports, AD diagnosis had a significant total effect on APP770 levels (*c* = 0.699, *p* = 3.1 × 10⁻⁴). This relationship was significantly mediated by TDP-43 pathology: AD diagnosis strongly predicted pTDP-43 levels (*a* = 0.073, *p* = 1.6 × 10⁻⁷), which in turn drove increased APP770 (*b* = 1.404, *p* = 0.025). The direct effect of AD diagnosis remained significant but markedly attenuated (*c’* = 0.597, *p* = 0.0026), suggesting that TDP-43 is a key mediator linking AD to pro-amyloidogenic *APP* splicing.

### Spatial transcriptomics of TDP-43-dependent *APP*/*STMN2* mis-splicing

We performed the first spatial transcriptomics study of LATE hippocampi, using the Xenium platform to relate phosphorylated TDP-43 (pTDP-43) pathology to splicing dysregulation. We designed a custom probe set targeting the APP695 and APP751/770 mRNA isoforms and the *STMN2* cryptic exons and analyzed hippocampal sections from LATE-NC stage 0 (no detectable pTDP-43 pathology) and stage 2/3 (extensive pTDP-43 pathology). Spatially quantified pTDP-43 burden was highly consistent with standard neuropathological classification (*P* < 0.01, **Figure 3A-C**). The *APP* isoform ratio 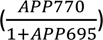 was significantly higher in spatial bins containing pTDP-43-positive cells than in pTDP-43-negative bins (*P* < 0.01, **Figure 3D**); reciprocally, pTDP-43 burden was higher in bins with high versus low *APP* ratios (*P* < 0.05, **Figure 3E**). This spatial correlation reinforces our bulk RNA-seq findings (ROSMAP and Kentucky cohorts) and sorted C9-ALS/FTD-TDP nuclei data ^29^, confirming that *APP* splicing abnormalities are physically linked to sites of TDP-43 proteinopathy.

**Figure 3:**
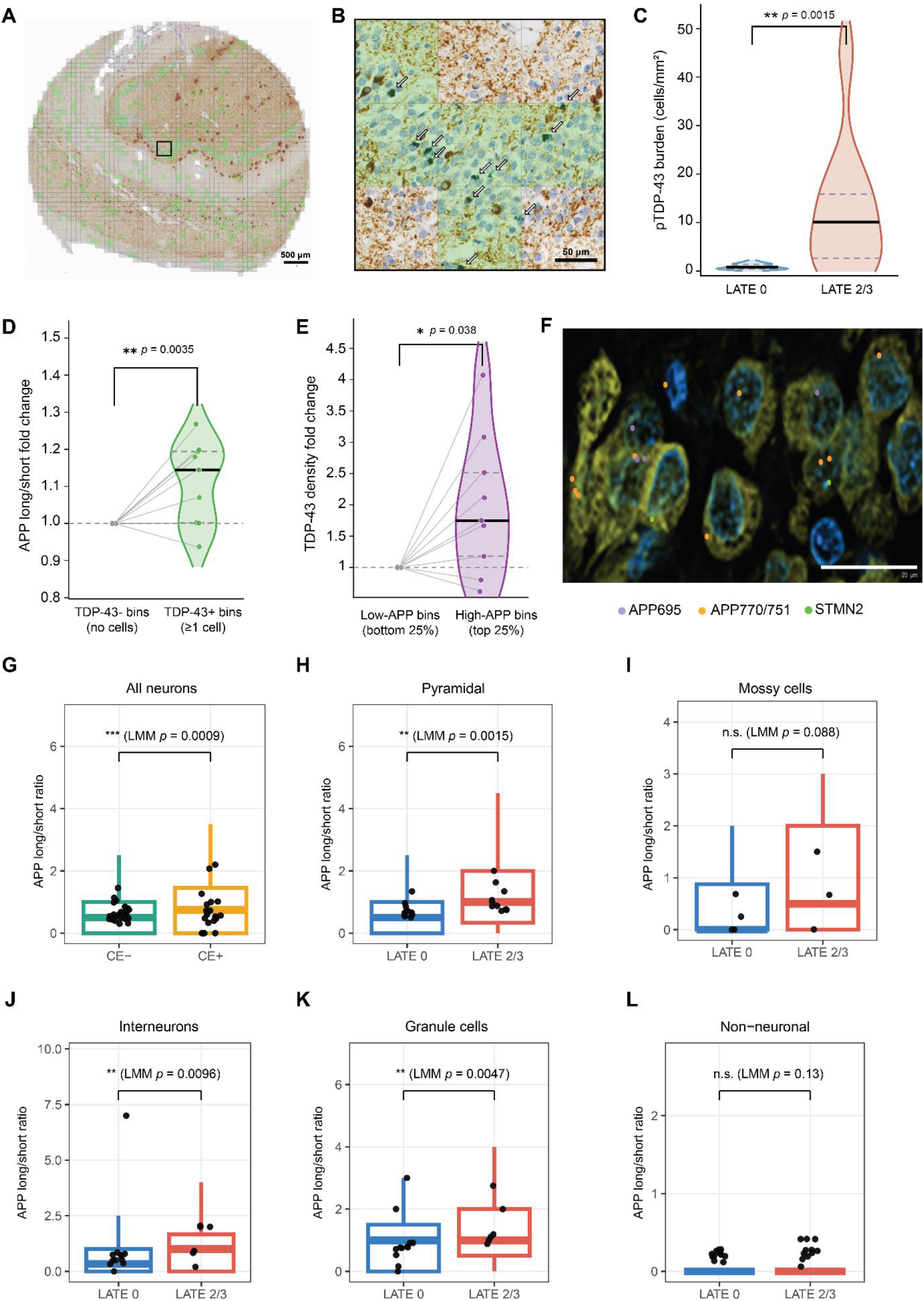
Spatial transcriptomic analysis of *APP* isoforms in LATE-NC hippocampal tissue. **(A)** Whole-section overview of LATE Stage 3 CA1 hippocampus, co-stained for pTDP-43 and Tau (green or brown chromogen, respectively). The section is partitioned into 50^2^ μm spatial bins, registered to paired Xenium transcriptomics data. Overlay: green, TDP-43+ bins; grey, TDP-43-bins; uncolored, non-tissue background. Scale bar, 500μm. **(B)** Magnified inset of the region indicated by the black rectangle in **(A)**, scale bar, 50μm. Arrows indicate individual pTDP-43+ cells identified by automated segmentation of the green chromogen signal. **(C)** pTDP-43 burden (cells/mm^2^) by LATE stage (LATE 0, *n* = 11; LATE 2/3, *n* = 10). **(D)** *APP* long/short isoform ratio 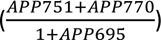, in TDP-43+ spatial bins relative to TDP-43-bins of the same donor, and reciprocally **(E)** fold change in pTDP-43 cell density between spatial bins in the top and bottom *APP* long/short isoform ratio quartiles, per donor. (C-E) Violin plots show median (solid line) and interquartile range (dashed lines). Two-sided Mann–Whitney U test; in D-E, per-donor values were compared with 1 (*n* = 9 LATE 2/3). **(F)** Xenium image representative of 24 donors (*n* = 12, 12; LATE 0, LATE 2/3) showing spatial distribution of APP695 (purple), APP751/770 (orange) and *STMN2* cryptic exon (CE, green) transcripts. Blue: nuclei; yellow: cytoplasm. Scale bar: 20 µm. **(G)** Ratio of long/short *APP* isoforms in all neurons classified by *STMN2* CE status (CE-, *n* = 13,163 cells from 23 donors; CE+, *n* = 563 cells from 21 donors) across all LATE stages. Comparison of long/short *APP* isoform ratios between LATE 0 and LATE 2/3, in **(H)** pyramidal neurons (*n* = 12, 10), **(I)** mossy cells (*n* = 4, 3), **(J)** interneurons (*n* = 12, 6), **(K)** granule cells (*n* = 12, 6), or **(L)** non-neuronal cells (*n* = 12, 11). Box plots of individual cells: center line, median; box bounds, 25th-75th percentiles; whiskers, the most extreme values within 1.5× the interquartile range of the box; outlying points not shown. Black dots, per-donor ratio. Linear mixed model with group as a fixed effect and donor as a random intercept; two-sided t-test with Satterthwaite’s degrees of freedom. Unless otherwise stated, *n* denotes biologically independent donors. No adjustment for multiple comparisons was made. Exact P values are shown in the figure. *P < 0.05, **P < 0.01, ***P < 0.001; n.s., not significant.

At the cellular level, neurons harboring *STMN2* cryptic exons displayed a significantly increased *APP* long-to-short isoform ratio (all neurons, *STMN2* CE− (n = 13,163) vs. CE+ (n = 563) across all LATE stages; *P* < 0.001, **Figure 3G**). This shift was also evident in pyramidal neurons (LATE 0 (n = 3,955) vs. LATE 2/3 (n = 1,695)), granule cells and interneurons (*P* < 0.001) of LATE 2/3 relative to LATE 0 specimens (**Figure 3H-K**), but not significant in non-neuronal cells (*P* > 0.05, **Figure 3L**). These data provide high-resolution spatial evidence associating TDP-43 pathology with *APP* and *STMN2* splicing dysregulation in the human hippocampus.

### TDP-43 toxic gain of function leads to *APP* mis-splicing

We next asked whether TDP-43 loss *per se* is associated with APP mis-splicing. Published and newly generated RNA-seq data of TDP-43 knockdown (KD) in iPSC-derived neurons ^10^ revealed no aberrant *APP* splicing (ΔPSI < 0.01, *P* > 0.6; **Figure 4A** and **Supplementary Table 3**). Similarly, no *APP* mis-splicing was observed upon complete TDP-43 knockout in HeLa cells ^31^ (ΔPSI < 0.01, *P* > 0.55, **Figure 4B** and **Supplementary Table 3**). Moreover, only 23% of mRNAs mis-spliced in TDP-43-negative ALS/FTLD-TDP brain nuclei were affected by TDP-43 KD (Fisher’s exact, *P* < 0.0001, **Figure 4C**). Thus, a substantial portion of splicing changes in postmortem tissue cannot be explained solely by loss of TDP-43 function and likely involves additional mechanisms.

**Figure 4:**
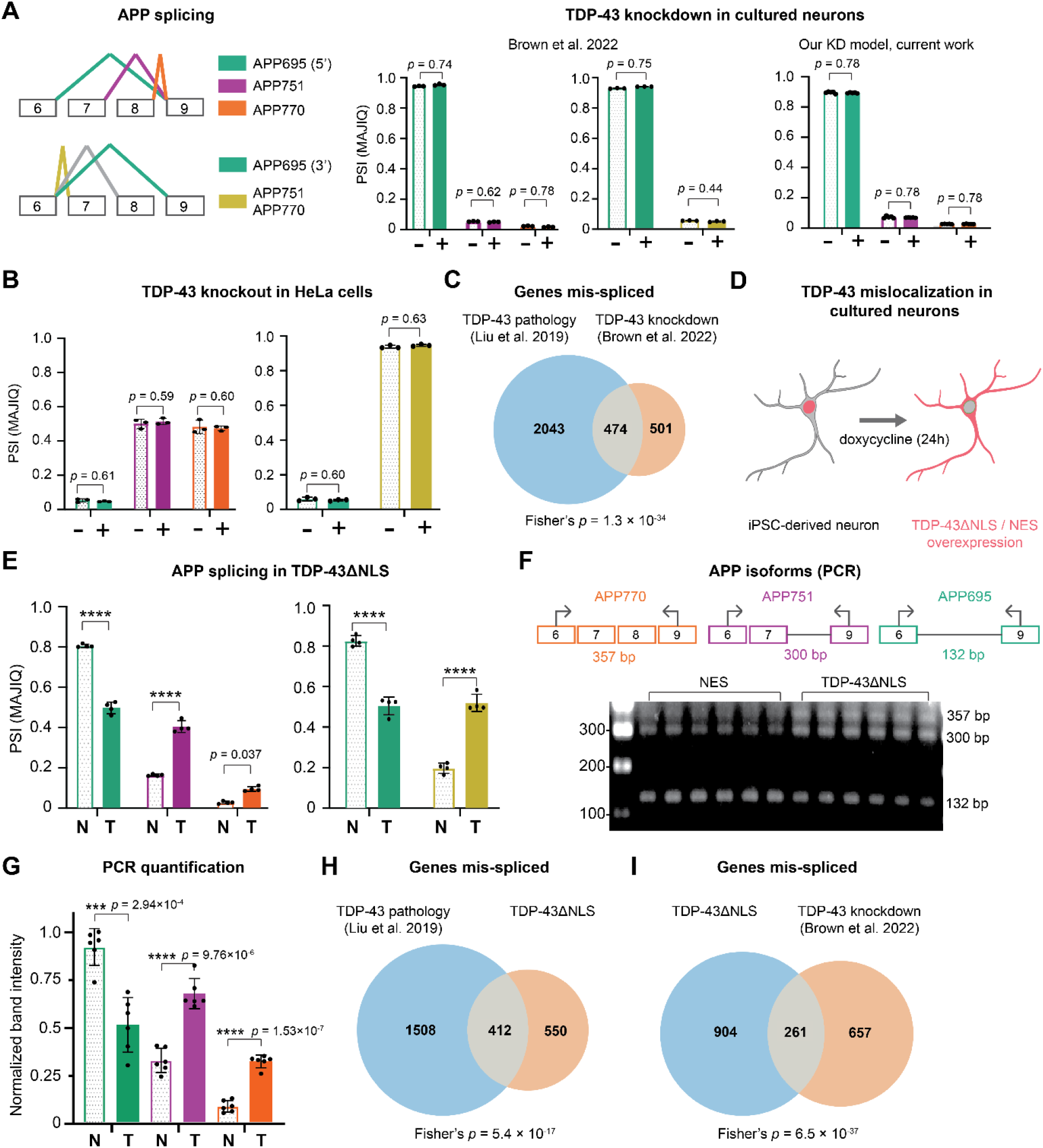
Gain of toxic function of cytoplasmic TDP-43 mislocalization causes *APP* mis-splicing. PSI of splice junctions corresponding to the three main *APP* isoforms upon TDP-43. **(A)** KD in cultured neurons from Brown et al. ^10^, (*n* = 3; our KD model, *n* = 4). The left/right bar plots correspond to the upper/lower junction blocks in the scheme, respectively. **(B)** TDP-43 knockout in HeLa cells ^31^ (*n* = 3; control (-); KO (+)). **(C)** Overlap of mis-spliced mRNAs in ALS/FTLD-TDP postmortem neuronal nuclei displaying TDP-43 pathology ^29^ and TDP-43 KD in cultured neurons ^10^. **(D)** Experimental setup: iPSC-derived cortical neurons ^74^, where NLS-mutated TDP-43 (TDP-43^ΔNLS^) or nuclear export signal (NES) expression was induced with doxycycline at day 8. **(E)** PSI of the same junctions in neurons expressing NES (N) or TDP-43ΔNLS (T); *n* = 4 independent differentiations per condition. PSI values were extracted from MAJIQ VOILA; conditions were compared with MAJIQ ΔPSI (*P* = 1 - posterior probability of |ΔΨ| > 0.02); **** indicates a MAJIQ *P* < 1 × 10⁻⁴, the smallest value MAJIQ reports. **(F)** RT-PCR of *APP* isoforms in NES or TDP-43^ΔNLS^ neurons, using primers spanning *APP* exons 6 to 9; samples were run on the same gel (uncropped gel in Supplementary Figure 1a). **(G)** Quantification of PCR band intensities, reflecting relative isoform abundance. *n* = 6 independent differentiations, arbitrary units. Two-sided Welch’s t-test. Overlap of mis-spliced mRNAs in **(H)** ALS/FTLD-TDP postmortem neuronal nuclei displaying TDP-43 pathology ^29^ versus TDP-43^ΔNLS^ neurons, and **(I)** TDP-43^ΔNLS^ neurons versus TDP-43 KD neurons ^10^. (C,H,I) Only mRNAs detected in both datasets were compared (|ΔPSI| > 0.1, *P* < 0.1); overlap exceeded chance expectation (one-sided Fisher’s exact test). Bar plots show mean ± s.d.; dots represent individual samples. No adjustment for multiple comparisons. \*\**P* < 0.01, \*\*\**P* < 0.001, \*\*\*\**P* < 0.0001; exact P values are shown in the panels.

We hypothesized that *APP* splicing is affected by cytoplasmic mislocalization of TDP-43, a core pathological feature that is absent after TDP-43 KD. To test this, we overexpressed TDP-43 carrying a mutated nuclear localization signal (NLS), rendering it cytoplasmic, in iPSC-derived neurons ^32^. Neurons overexpressing a nuclear export signal (NES) served as a cytoplasmic overexpression control (**Figure 4D**). RNA-seq revealed 32% lower usage of the APP695-specific junction in TDP-43^ΔNLS^ relative to NES (*P* < 0.0001, **Figure 4E** and **Supplementary Table 3**). In contrast, usage of the APP751-specific junction and of the exon 6-7 junction increased by 24% and 32%, respectively (*P* < 0.0001, **Figure 4E**). Overall APP levels were reduced twofold (*P* < 0.0001, **Supplementary Table 5**). Reverse transcription PCR of the three major isoforms (**Figure 4F; Supplementary Figure 1a**) verified a 19% reduction in APP695 (*P* < 0.01) and a 4-fold increase in APP770 (*P* < 0.0001) in TDP-43ΔNLS iPSC-derived neurons (**Figure 4G**), consistent with *APP* mis-splicing in ALS/FTLD-TDP and AD brains with TDP-43 pathology. More broadly, about 27% of mRNAs mis-spliced with TDP-43 pathology in ALS/FTLD-TDP tissue ^29^ were also mis-spliced in TDP-43^ΔNLS^ neurons (Fisher’s exact, *P* < 0.0001, **Figure 4H**). This is comparable to TDP-43 KD, suggesting that toxic gain-of-function through mislocalization similarly contributes to splicing defects in ALS/FTLD-TDP. About 25% of splicing defects in the Fratta lab TDP-43 KD model ^10^ were recapitulated in TDP-43^ΔNLS^ neurons (Fisher’s exact, *P* < 0.0001; **Figure 4I**). However, ΔPSI concordance across the 238 shared junctions was significantly lower than the Fratta lab TDP-43 KD ^10^ and our TDP-43 KD (**Extended Data Figure 3A,B**). Moreover, 59 genes were mis-spliced in both TDP-43 KD and TDP-43^ΔNLS^ models, an overlap significantly greater than expected by chance (Fisher’s exact, *P* < 0.0001), but these did not include *STMN2* or *UNC13A*. Together, these findings identify TDP-43 mislocalization as a toxic gain-of-function mechanism for mis-splicing, distinct from TDP-43 loss-of-function.

### TDP-43 drives RBPs cytoplasmic mislocalization

As many ALS/FTLD-TDP splicing abnormalities could not be explained solely by nuclear loss of TDP-43, we hypothesized that additional splicing regulators are mislocalized with TDP-43 to the cytoplasm. To test this, we performed nucleo-cytoplasmic fractionation and mass spectrometry of induced neurons expressing TDP-43^ΔNLS^ or NES (**Extended Data Figure 4A** and **Supplementary Table 6**). The nuclear proteins Lamin-B1 and HDAC1 were highly enriched in the nuclear fraction of both TDP-43^ΔNLS^ and control NES lines (*log2FC* = 5.7 / 5.1, *FDR* = 0; **Extended Data Figure 4B**), and gene ontology analysis of the top 200 proteins most enriched in or depleted from the nucleus ^33,34^ confirmed proper separation of fractions (**Extended Data Figure 4C,D**). TDP-43^ΔNLS^ expression was associated with cytoplasmic enrichment of 173 proteins relative to NES control neurons (*FDR* < 0.05, *log2FC* > 1; **Extended Data Figure 4E**). TDP-43 itself was among the most enriched (6.5-fold; *FDR* = 0.017), and 20 of the mislocalized proteins are known splicing regulators: CELF1, CPSF7, CRIPT, DDX17, DHX16, ERH, FAM98B, HNRNPLL, LGALS3, NONO, PSPC1, PTBP2, SNRPC, SRSF1, SRSF2, SRSF4, SRSF5, SRSF6, SUPT6H and U2SURP. TDP-43^ΔNLS^ increased the cytoplasmic-to-nuclear ratio of these 20 splicing regulators relative to NES (**Extended Data Figure 4F**), suggesting that additional splicing regulators may be mislocalized to the cytoplasm together with TDP-43 in human neurons.

To elucidate the mechanisms of aberrant splicing downstream of cytoplasmic TDP-43, we further characterized proteins in its vicinity, leveraging our recent proximity labeling proteomics dataset from TDP-43^ΔNLS^-APEX neurons ^32^ (**Extended Data Figure 4G-I**). Upon hydrogen peroxide (H2O2) exposure, APEX generates biotin-phenol radicals that label neighboring biomolecules. To probe cytoplasmic TDP-43 surroundings within stress granules (cytoplasmic biomolecular condensates), we also applied sodium arsenite (250 µM, 90 minutes) in two independent biological experiments.

TDP-43 itself was among the most enriched proteins (*FDR* = 0, *log2FC* = 5.24; red dots in **Extended Data Figure 4H,I**). Although total protein numbers differed between experiments, 212 proteins were detected in both independent repeats under basal (unstressed) conditions and 229 under stress (*FDR* < 0.01, *log2FC* > 2). 126 proteins were consistently associated with cytoplasmic TDP-43 regardless of stress state (**Extended Data Figure 4G)** ^32^. Gene Ontology (GO) biological process analysis revealed enrichment of pathways regulating gene silencing, cell cycle, transcription and translation (**Extended Data Figure 4J-K**), in line with known TDP-43 functions ^35–39^. The TDP-43-associated proteins included 11 splicing regulators: 3 under basal conditions only (CELF2 ^40^, METTL3 ^41^ and TIAL1 ^42^), 5 under stress only (CASC3 ^43,44^, ECD ^45,46^, IVNS1ABP ^47^, TTF2 ^48^ and ZCCHC8 ^49^), and 3 under both (SCAF11 ^50^, TARDBP ^8^ and TIA1 ^51,52^).

### Loss of splicing regulators mimics TDP-43 failure

We next tested whether the newly identified TDP-43 interactors regulate splicing by generating loss-of-function (knockdown, KD) alleles with a cytidine base editor ^53^ in human iPSCs differentiated into cortical neurons. In total, the fractionation and proximity labeling experiments identified 31 splicing regulators. As the two approaches did not clearly overlap, we shortlisted 8 proteins based on both experiments and prior literature: CELF2 ^54^, CPSF7 ^55^, HNRNPLL ^56^, SCAF11 ^50^, SRSF5 ^57^, SUPT6H ^58^, TIA1 ^59^ and TIAL1 ^59^ (**Figure 5A**). TIA1 was included for its genetic association with ALS ^60,61^, with its paralog TIAL1. Premature stop codons reduced mRNA levels and plausibly prevented full-length protein production (**Extended Data Figure 5; Supplementary Figure 1b**). RNA-seq splicing analysis revealed differential usage (|ΔPSI| > 0.1, *P* < 0.1) of hundreds of splice junctions upon KD of the eight regulators (**Extended Data Table 1** and **Supplementary Table 2**). TIA1 displayed the mildest phenotype, likely because only a heterozygous KD could be generated (**Extended Data Table 1**).

**Figure 5:**
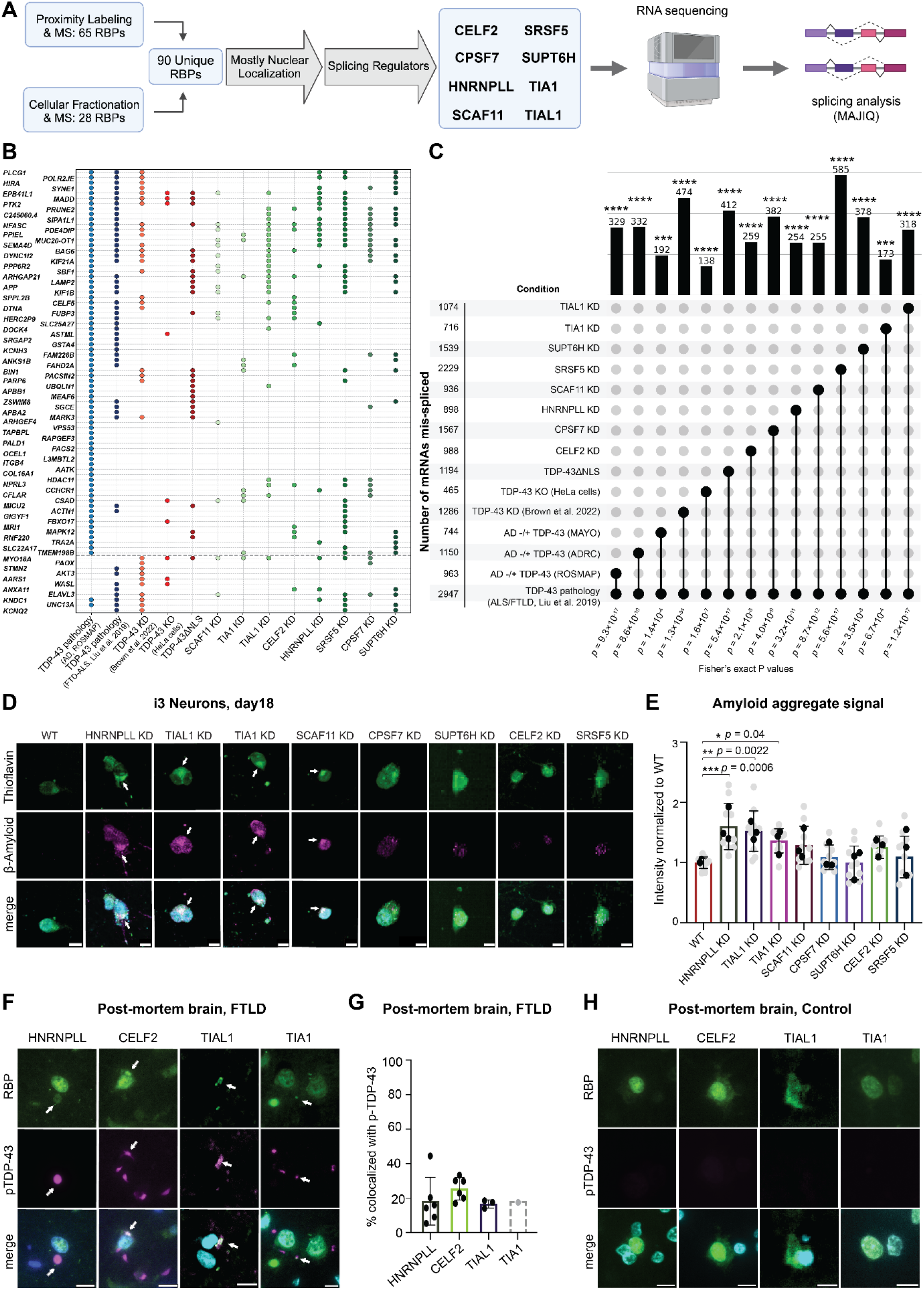
TDP-43 partner RBPs partially mimic disease splicing changes.(A) Prioritization of splicing regulators for in-depth molecular and neuropathological investigation. **(B)** Bubble heat map of junction-level |ΔPSI| for the 73 high-confidence mRNAs affected by TDP-43 proteinopathy in AD (Fig. 1J) and 11 mRNAs affected by TDP-43 loss of function, across postmortem datasets and cellular models. **(C)** Upset plot of mis-spliced mRNAs per condition (rows) and those shared between datasets and C9-ALS/FTD-TDP pathology ^29^ (columns). Thresholds: |ΔPSI| > 0.1, *P* < 0.1 (cell models); |ΔPSI| > 0.02, *P* < 0.05 (patient data). Only mRNAs detected in both datasets were compared. One-sided Fisher’s exact test. **(D)** Representative images of Aβ (anti-Aβ1-16, magenta) co-localized with Thioflavin S aggregate (green) on day 18 KOLF2.1J (wild-type, WT) iPSC-derived neurons with specific RBP knockdown. Arrows - co-localization; scale bars, 10 µm. Images are representative of *n* = 3 independent experiments with similar results. **(E)** Quantification of Aβ mean immunofluorescence intensity within Thioflavin S aggregates, normalized to WT. *n* = 3 independent differentiations (black dots) and technical replicates (grey dots, 98 images/repeat). Mean ± s.d. One-way ANOVA with two-sided Dunnett’s multiple-comparisons test versus WT. **(F)** Representative images of pTDP-43 and RBP immunohistofluorescence in C9-ALS/FTD-TDP brains; arrows - extranuclear co-localization. **(G)** Percentage of RBP co-localizing with pTDP-43 inclusions, per case. FFPE sections for HNRNPLL (*n*=6), CELF2/CUGBP2 (*n*=6) and TIA1 (the dashed bar denotes reduced confidence inherent to a single positive observation in 6 cases). TIAL1 (cryosections, *n* = 3). Mean ± s.d.; **(H)** Representative immunostaining in non-neurological control postmortem tissue. In F and H, images are representative of *n* = 6 (HNRNPLL, CELF2/CUGBP2, TIA1) and n = 3 (TIAL1) biologically independent cases with similar results; scale bars, 5 µm. *P < 0.05, **P < 0.01, ***P < 0.001, ****P < 0.0001; exact P values are shown in the panels.

We next compared splicing at previously reported TDP-43-dependent junctions across published data ^10,20,29,31,62,63^, RNA-seq of neurons with KD of the eight splicing regulators, and control induced neurons. Cryptic exon splicing in *STMN2* and *UNC13A* was detected upon TDP-43 KD and with TDP-43 pathology in ALS/FTLD-TDP and AD tissues (**Figure 5B**). *UNC13A* was also mis-spliced upon SRSF5 and SUPT6H KD, but at junctions different from those previously reported. In SRSF5 KD neurons, two UNC13A events were significantly altered: one between canonical exon 7 and an unannotated exon identified by MAJIQ (hg38; chr19:17665549-17665977, |ΔPSI| = 17%, *p* = 0.057), and another involving exon 37 and canonical exon 39 (hg38; chr19:17623118-17623263, |ΔPSI| = 20%, *p* = 0.018) (**Extended Data Figure 6A**). The same mis-splicing between exon 7 and an unannotated exon occurred upon SUPT6H KD (|ΔPSI| = 18%, *p* = 0.035, **Extended Data Figure 6B**). Each event was relatively rare, with only a few dozen reads per junction. Reported mis-splicing events in *MYO18A* and *ANXA11* were also found upon KD of several of these regulators (**Figure 5B**). *ANXA11* and *APP* were mis-spliced in postmortem tissues and upon RBP KD, but not upon TDP-43 KD, suggesting that nuclear loss of these regulators, rather than of TDP-43 itself, causes their aberrant splicing.

More broadly, KD of the eight splicing regulators consistently recapitulated dozens, and in several instances hundreds, of splicing events associated with TDP-43 pathology in C9-ALS/FTLD, AD DLPFC ^20,29^ or TDP-43^ΔNLS^ (**Figure 5C, Extended Data Figure 7A,B**, Fisher’s exact *P-values* < 0.001, < 0.01, < 0.0001, respectively). For example, 30% or 20% of mRNAs mis-spliced in ALS/FTLD-TDP CNS were also mis-spliced upon TIAL1 or SRSF5 KD (318 of 1074, or 585 of 2947, respectively, Fisher’s exact *P* < 0.0001, **Figure 5C**). Splicing alterations were concordant between patient brain RNA-seq and RBP KD models, and between TDP-43^ΔNLS^ and both the brain RNA-seq and RBP KD datasets. By contrast, TDP-43 KD-associated changes were concordant with only a subset of the pathologies or RBP KD models (**Extended Data Figure 7C**). These results suggest that cytoplasmic TDP-43 mislocalization induces splicing defects through both nuclear loss-of-function and interference with other splicing regulators, underscoring the relevance of the TDP-43^ΔNLS^ model as a surrogate for TDP-43 proteinopathy.

### Splicing regulators control *APP* mis-splicing and Aβ

We next examined whether loss of TDP-43-associated splicing regulators could cause the *APP* mis-splicing seen upon TDP-43 mislocalization. APP770 levels increased upon KD of TIAL1, SRSF5, SCAF11 or CELF2 (*APP* exon 8-specific PCR band, *P* < 0.05, **Extended Data Figure 7F; Supplementary Figure 1c**), without significant change in global *APP* mRNA levels in the RBP KD lines (**Supplementary Table 5**). TIAL1, SCAF11 or SRSF5 KD also increased usage of the *APP* exon 7-8 junction (*P* < 0.0001, **Extended Data Figure 7D** and **Supplementary Table 3**). Exon 7-9 junction usage increased upon TIAL1 KD (*P* < 0.05), and exon 8-9 junction usage upon TIAL1 and SRSF5 KD (*P* < 0.05, **Extended Data Figure 7E** and **Supplementary Table 3**). These results suggest that nuclear depletion of splicing regulators through co-sequestration with TDP-43 shifts the *APP* isoform balance towards APP751 and APP770. Moreover, five of the candidate factors bind *APP* RNA according to the POSTAR3 database ^64^: TIA1 and TIAL1 (eCLIP), CPSF7, SRSF1 (PAR-CLIP), and CELF2 (HITS-CLIP).

Finally, thioflavin-positive Aβ aggregate intensity increased in HNRNPLL, TIAL1 and TIA1 KD neurons relative to control (**Figure 5D,E**) and knockdown of HNRNPLL, TIAL1 and TIA1 or the TDP-43^ΔNLS^ model, further increased pTau aggregate intensity, *in vitro* (**Extended Data Figure 8**). However, co-localization of RBPs with pTau was not detected in AD/LATE postmortem tissue, pointing to potential context-dependent differences between acute RBP depletion and terminal neuropathology.

### Splicing regulators in FTLD-TDP postmortem inclusions

To validate sequestration of splicing regulators by TDP-43, we examined RBP colocalization in C9-ALS/FTD-TDP neuropathological specimens. HNRNPLL, CELF2/CUGBP2 and TIAL1 were found within cytoplasmic pTDP-43 inclusions, colocalizing in 5-44% of analyzed cells across cases (**Figure 5F-H)**. Rare TIA1/pTDP-43 colocalization (1 of 6 cases) aligns with prior reports ^65,66^. SRSF5 showed no detectable colocalization, and suitable antibodies for SCAF11 and SUPT6H were lacking. However, in TDP-43^ΔNLS^ induced neurons, we identified cytoplasmic mislocalization of TIAL1, SCAF11 and SRSF5 (**Extended Data Figure 9A-D**). These findings confirm that a subset of splicing regulators is directly recruited into neuropathological pTDP-43 inclusions.

## Discussion

Despite the established role of splicing dysregulation in neurodegeneration, including AD, its upstream drivers remain poorly understood. TDP-43 proteinopathy is strongly linked with ALS, FTD, AD and LATE, and abnormal TDP-43 state contributes to cell loss.

Here, we demonstrate that TDP-43 pathology is a significant driver of splicing abnormalities in AD, explaining about 20% of AD splicing changes. This far exceeds any previously reported mechanism of splicing regulation in AD, establishing TDP-43 dysfunction and impaired splicing as unifying features across multiple neurodegenerative diseases. TDP-43 splicing pathology is evident in the hippocampus and amygdala, where LATE neuropathological changes are well characterized. The unexpected, consistent splicing dysregulation in AD prefrontal cortex, despite low TDP-43 burden ^2,13,14^, may reflect subtle functional changes undetectable by pTDP-43 immunohistochemistry, or axonal transport of cryptic mRNAs from the hippocampus or amygdala to the prefrontal cortex.

*APP* alternative splicing has been recognized in AD for decades, but its mechanism remained unclear. Our study provides insight into the regulation of *APP* splicing downstream of dysregulated TDP-43 in ALS/FTLD-TDP and AD. In this first spatial transcriptomic study of hippocampus with LATE-NC, *APP* splicing dysregulation correlated with pTDP-43 pathology and *STMN2* cryptic exon inclusion, was specific to advanced LATE stages and spanned several neuronal cell types. TDP-43 regulation of *APP* splicing is substantiated by sorted-nuclei transcriptomics from *C9orf72* ALS/FTD brains, by mechanistic validation in TDP-43^ΔNLS^ iPSC-derived neurons and by mediation analysis, which reveals that TDP-43 pathology is associated with Aβ burden both directly and indirectly via a shift toward longer *APP* isoforms. This aligns with findings where longer APP isoforms increased Aβ production *in vitro* ^67^, AD mouse models where hippocampal TDP-43 increased plaques and impaired memory ^68^, and human data correlating KPI(+)APP with plaque density ^69^. The TDP-43–Aβ link also suggests an underexplored contribution to ALS-FTD pathogenesis.

Driven by nuclear depletion, TDP-43 loss-of-function triggers cryptic exon splicing ^5,9–12^, a phenomenon documented in AD ^16–18^. Beyond loss-of-function, we demonstrate a non-mutually exclusive toxic cytoplasmic gain-of-function, in which TDP-43 sequesters splicing-related RBPs, indirectly driving splicing pathology. We provide evidence that the RBPs CPSF7, SCAF11, SUPT6H, SRSF5, TIAL1, TIA1, CELF2 and HNRNPLL contribute to disease-relevant splicing in AD, and the latter four colocalize with pTDP-43 inclusions. *TIA1* genetic variants are associated with ALS/FTD, and TIA1 protein colocalizes with TDP-43 in human ALS spinal cord tissue and cultured cells ^60^.

Loss of SRSF5, SCAF11 or TIAL1 alters *APP* splicing, and TIAL1 also binds *APP* mRNA ^64^; thus, TIAL1, and perhaps SRSF5 and SCAF11, may mediate the effect of TDP-43 on APP alternative splicing. However, of these three, only TIAL1 colocalized with pTDP-43 inclusions, and none of the RBPs were enriched in the insoluble ALS/FTD brain proteome ^70^. These intersections warrant further functional exploration.

Given the prevalence of TDP-43 pathology in both LATE and AD, driving widespread RNA splicing errors and distorting protein output, this mechanism could fundamentally reshape our understanding of Alzheimer’s disease. Loss- and gain-of-function mechanisms cooperatively drive splicing dysregulation, indicating that TDP-43-targeted therapies must both restore nuclear function and mitigate cytoplasmic toxicity to be effective. Mirroring splice-correction strategies in SMA and ALS, modulating *APP* isoform splicing may offer a viable therapeutic avenue for AD.

Limitations: A substantial portion of AD splicing changes is not explained by TDP-43 dysregulation; other mechanisms remain uncovered. Although we identify cytoplasmic splicing regulators co-sequestered with TDP-43, their direct contributions to mis-splicing require further exploration, and RBPs not identified here may also contribute. Our proteomic analyses identified peptides specific to the longer APP isoforms but not an APP695-specific peptide, which would enable more complete protein-level analysis of isoform ratios. The mechanism through which longer *APP* isoforms increase Aβ pathology also remains to be investigated.

Our study suggests that TDP-43 pathology is a unifying molecular feature linking seemingly disparate neurodegenerative disorders and demonstrates that TDP-43 mislocalization leads to cytoplasmic sequestration of splicing regulators. By establishing TDP-43 as an upstream regulator of dozens of critical mRNAs, including *APP*, in ALS/FTD and AD, and as a contributor to amyloid pathology, we offer new insights into molecular convergence in neurodegeneration. Whether this convergence occurs early or is a late common consequence of diverse causes of chronic CNS neurodegeneration remain unknown. Targeting the TDP-43 pathway may offer therapeutic opportunities for ALS/FTLD-TDP and AD.

---

**Extended Data Figure 1:**
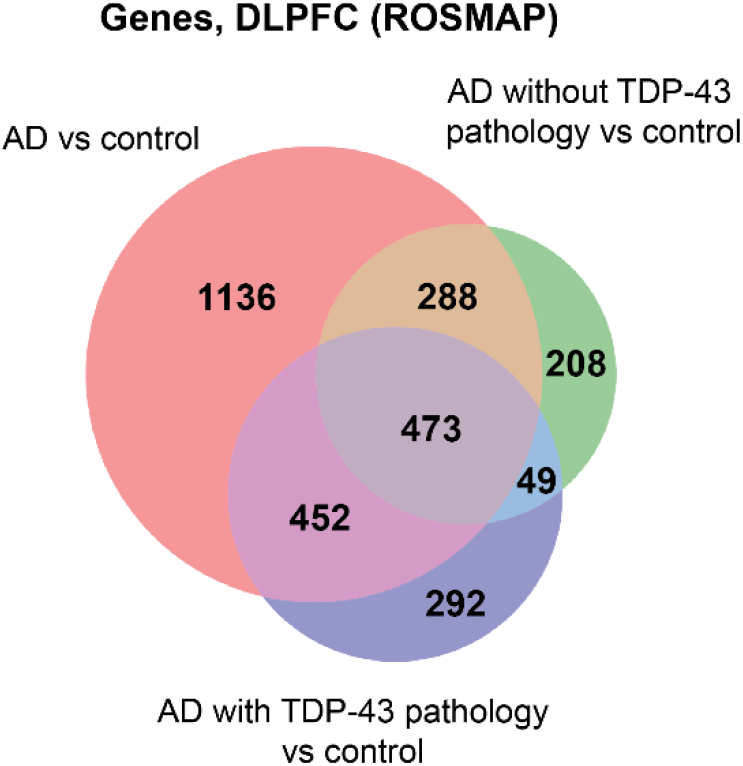
Splicing aberrations downstream of TDP-43 pathology in human postmortem tissue. Venn diagram illustrating the contribution of TDP-43 pathology to transcriptome-wide aberrant splicing changes in Alzheimer’s disease bulk RNA-seq. DLPFC from the ROSMAP cohort ^20^, comparing Alzheimer’s disease (AD) cases versus controls (n = 236, 200; red), AD cases without TDP-43 pathology versus controls (n = 126, 200; green) and AD cases with TDP-43 pathology versus controls (n = 110, 200; purple). Numbers reflect genes with any mis-splicing event (|ΔPSI| > 0.02, P < 0.05). Splicing was analyzed with MAJIQ Heterogen; P values are from the two-sided TNOM test, not adjusted for multiple comparisons.

**Extended Data Figure 2:**
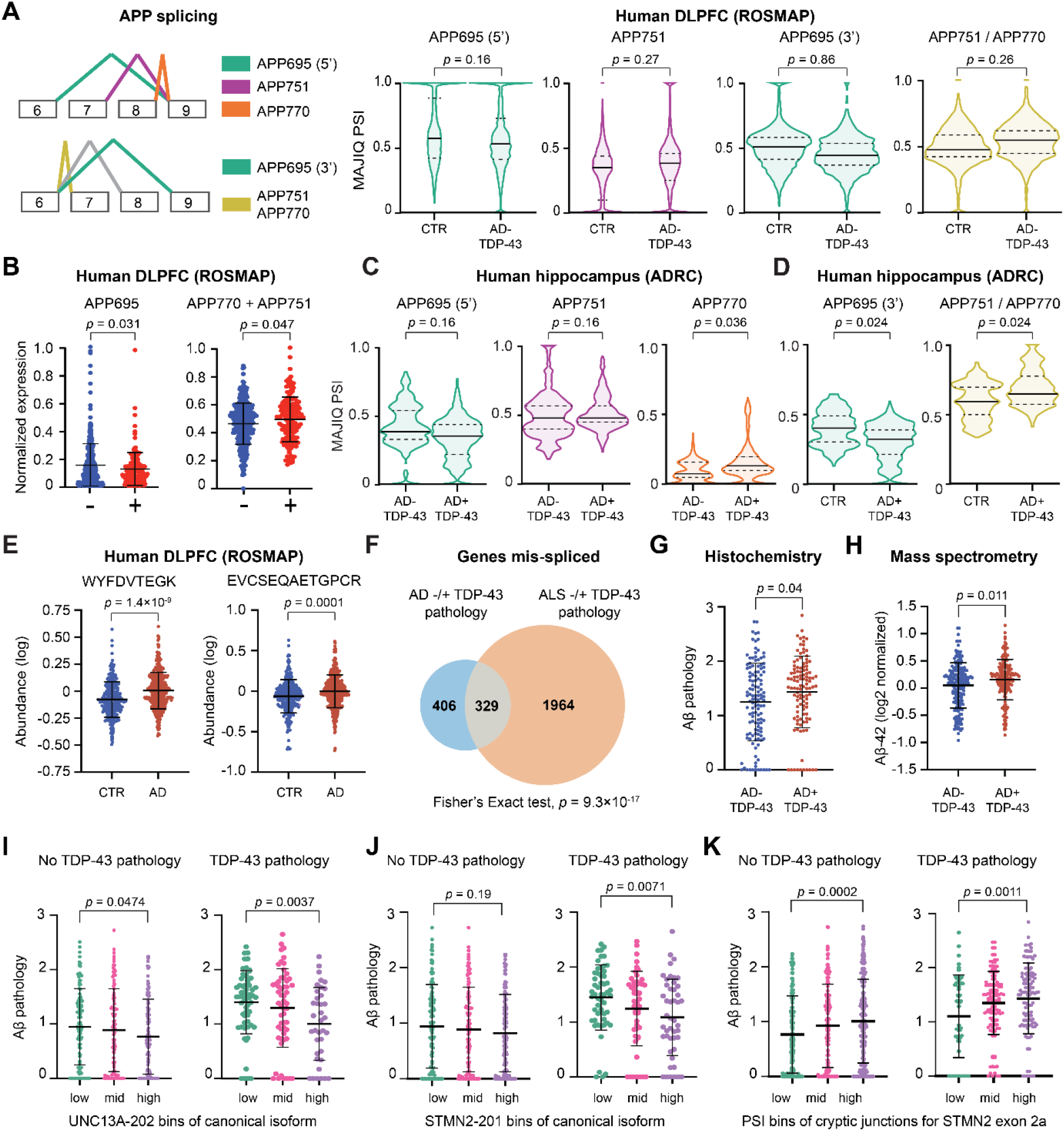
Aberrant splicing of *APP* downstream of TDP-43 pathology in human postmortem tissue. **(A)** MAJIQ splice junctions in APP exons 6-9 and corresponding isoforms (upper and lower junction blocks correspond to the first two and last two violin plots, respectively), with PSI values in ROSMAP DLPFC comparing controls with AD without TDP-43 pathology (*n* = 200, 126). **(B)** RSEM-estimated expression of APP695 (left) and APP751 + APP770 (right) in ROSMAP DLPFC samples without and with TDP-43 pathology (*n* = 349, 153); isoforms were min–max normalized and, for APP751 + APP770, summed and rescaled to 0–1. MAJIQ PSI values of *APP* exons 6-9 in ADRC hippocampus ^23^, comparing **(C)** AD+TDP-43 versus AD-TDP-43 (*n* = 35, 25) and **(D)** AD+TDP-43 versus controls (*n* = 35, 21; hippocampus cases ^23^ with pTDP-43 are classified as ‘LATE-NC + ADNC’ and those without as ‘ADNC’ or ‘AD’). In A, C and D, violin plots show the distribution of PSI values, with the median (solid line) and interquartile range (dashed lines); two-sided TNOM test implemented in MAJIQ Heterogen. **(E)** Abundance of two peptides mapping specifically to APP751 and APP770 comparing ROSMAP DLPFC from AD patients (*n* = 212) to cognitively unimpaired controls (n = 180). **(F)** Overlap of mRNAs mis-spliced in TDP-43-depleted vs TDP-43-positive nuclei from C9-ALS postmortem cortical tissue ^29^ and upon TDP-43 pathology in ROSMAP DLPFC AD samples; one-sided Fisher’s exact test. Aβ pathology in ROSMAP DLPFC samples comparing AD+TDP-43 (+) versus AD-TDP-43 (-) as measured by **(G)** immunohistochemistry ^73,78^ (*n* = 110, 125) or **(H)** mass spectrometry ^28^ (*n* = 178, 186; data were harmonized, batch-corrected and log2-transformed with TAMPOR). Aβ pathology (immunohistochemistry) in ROSMAP DLPFC samples binned into low, medium or high levels of **(I)** canonical *UNC13A* isoform (*UNC13A-202*, RSEM), **(J)** canonical *STMN2* isoform (*STMN2-201,* RSEM) or **(K)** MAJIQ PSI of the *STMN2* cryptic exon 2a junction, shown separately for samples without and with TDP-43 pathology (*n* = 349, 153). In B, E and G-K, data are presented as mean ± s.d. and groups were compared with a two-sided Welch’s t-test (in I-K, low versus high bin). No adjustment for multiple comparisons was made. Exact P values are shown in the figure.

**Extended Data Figure 3:**
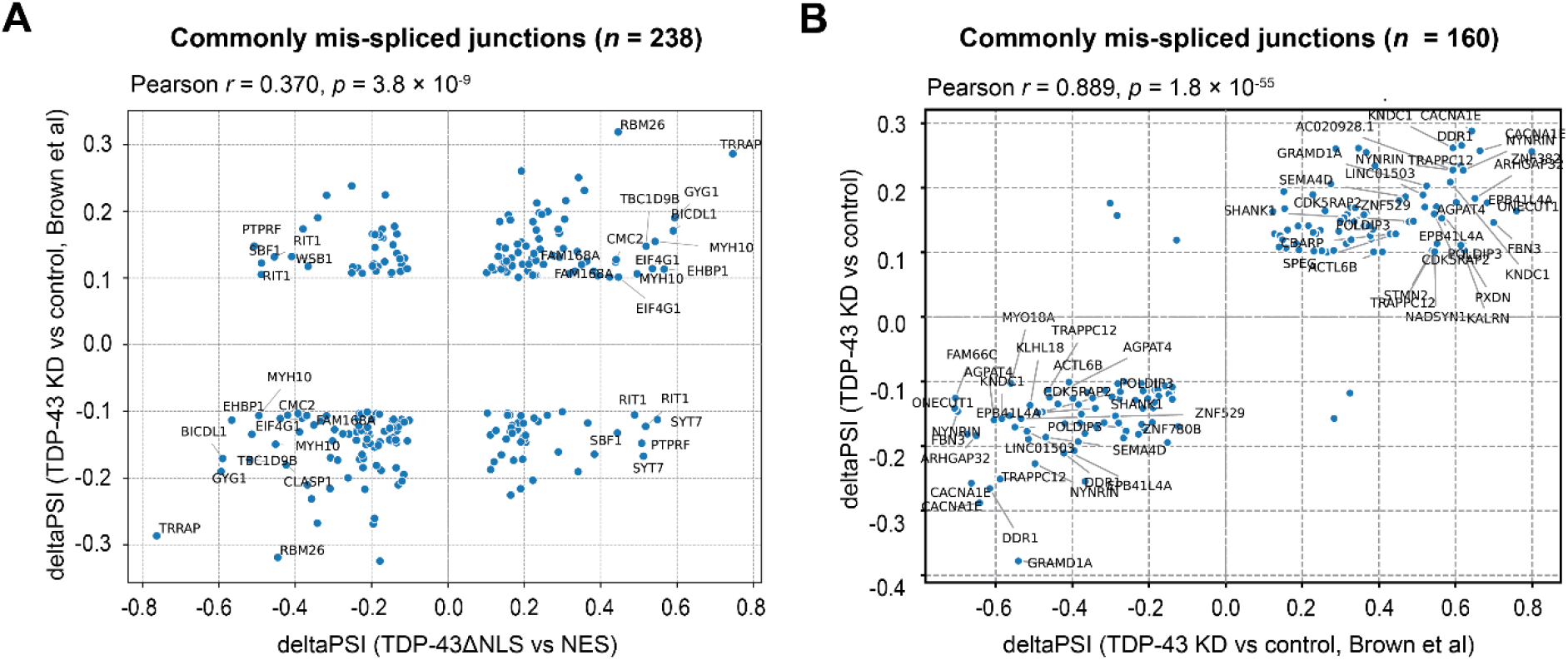
Concordance of splicing changes in TDP-43 knockdown and TDP-43^ΔNLS^ models. Scatter plot of splice junction-level percent spliced in (ΔPSI) changes: **(A)** TDP-43^ΔNLS^ relative to NES (x-axis) versus TDP-43 knockdown (data from ^10^) relative to control (y-axis), and **(B)** TDP-43 knockdown (data from ^10^) relative to control (x-axis) versus TDP-43 knockdown (CRISPRi, this study) relative to control. Junctions significantly mis-spliced in both datasets are shown (|ΔPSI| > 0.1, probability of change > 0.9; n = 238 and 160 junctions, respectively). mRNAs with |ΔPSI| > 0.4 in either comparison are labeled. Two-sided Pearson’s correlation, *r* and exact P values are shown in the panels.

**Extended Data Figure 4:**
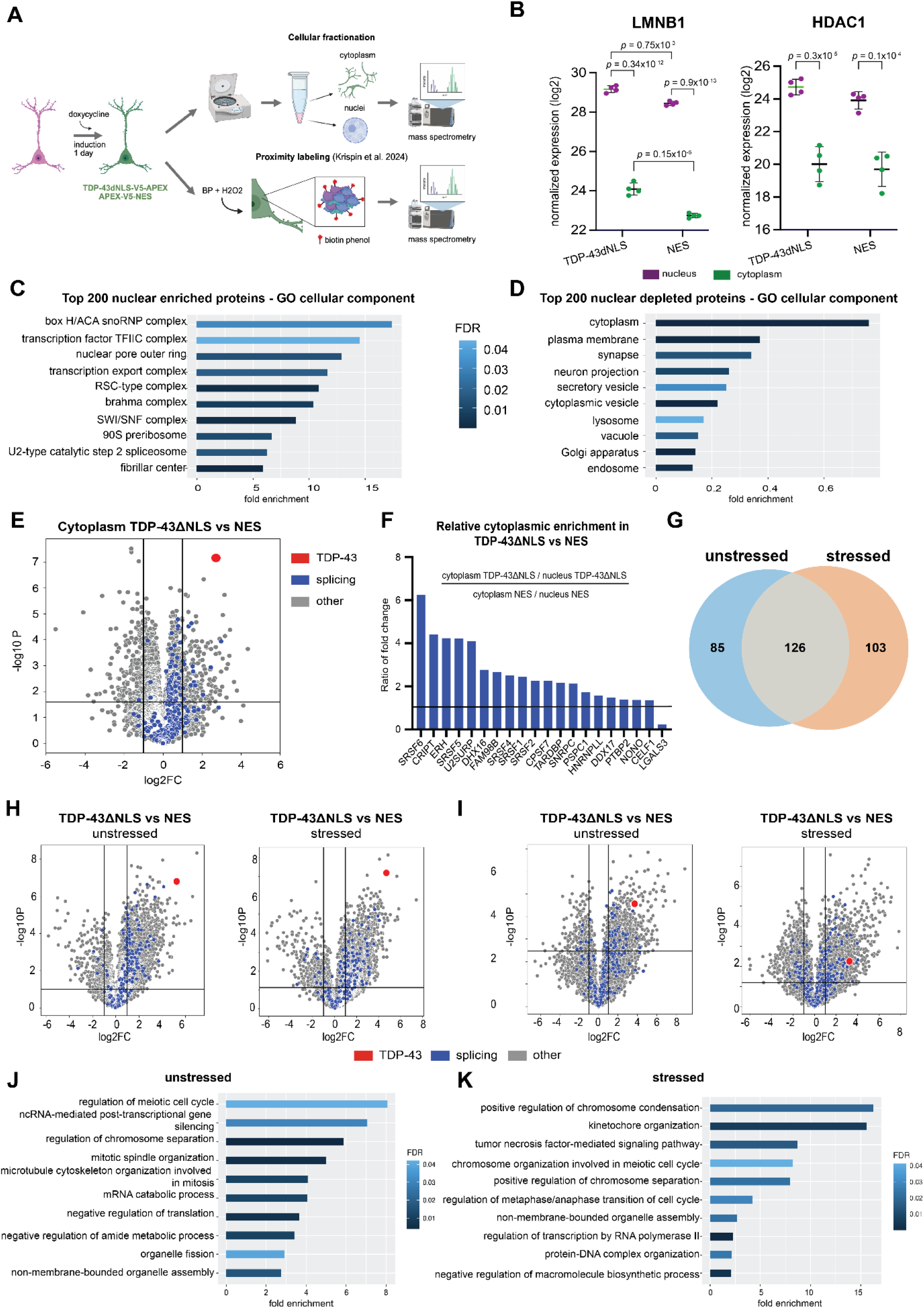
Probing cytoplasmic co-sequestration of splicing regulators by TDP-43. **(A)** Diagram of TDP-43^ΔNLS^ and cytoplasmic control construct (NES) expressed in cultured neurons for 1 day prior to harvest for unbiased proteomic analysis by mass spectrometry of nuclear and cytoplasmic fractions or by proximity labeling ^32^. **(B)** Relative LMNB1 and HDAC1 protein intensity in each of the subcellular fractions in either NES-or TDP-43^ΔNLS^-expressing neurons (*n* = 4 independent differentiations per condition). Values are log2-transformed expression values obtained with proteomics as described in the *Methods.* Two-way ANOVA, mean ± s.d. Gene ontology ^33,34^ cellular component analysis of the top 200 most nuclear **(C)** enriched or **(D)** depleted proteins. **(E)** Volcano plot showing differential protein abundance in the cytoplasm of neurons expressing TDP-43^ΔNLS^ compared with NES. Proteins annotated with the Gene Ontology term “splicing” are shown in blue; the protein TDP-43 is shown in red. Log2FC and P-values: two-sided t-test using log2-transformed MaxQuant LFQ values comparing TDP-43^ΔNLS^ (n=4) to NES (n=4) cytoplasmic fractions. P-values are adjusted (Perseus ^104^ q-value). Vertical lines indicate log2FC thresholds of −1 and 1; the horizontal line indicates an adjusted p-value threshold of 0.01. **(F)** Cytoplasmic enrichment of 20 splicing regulators in human neurons expressing TDP-43^ΔNLS^ relative to NES. Quantified cytoplasmic shift induced in human neurons expressing TDP-43^ΔNLS^ calculated by normalizing the cytoplasmic-to-nuclear ratio in TDP-43^ΔNLS^ to that in NES. **(G)** Venn diagram of candidate TDP-43 interactors identified reproducibly across two independent experiments, under unstressed and/or stressed conditions. Volcano plots of differential protein enrichment in TDP-43ΔNLS versus NES samples under basal conditions (left) or after sodium arsenite stress (right), in two independent biological experiments **(H)** experiment 1; **(I)** experiment 2, each with 3 or 4 technical repeats per condition. TDP-43 is shown in red, and proteins annotated with the Gene Ontology term ‘splicing’ in blue. Log2FC and P-values: two-sided t-test using log2-transformed MaxQuant LFQ values. P-values are adjusted (Perseus ^104^ q-value). Vertical lines indicate log2FC thresholds of −1 and 1; horizontal lines indicate an adjusted p-value threshold of 0.01. Gene ontology ^33,34^ “biological process” analysis of proteins enriched in TDP-43^ΔNLS^ samples versus NES samples in **(J)** unstressed or **(K)** stressed conditions.

**Extended Data Figure 5:**
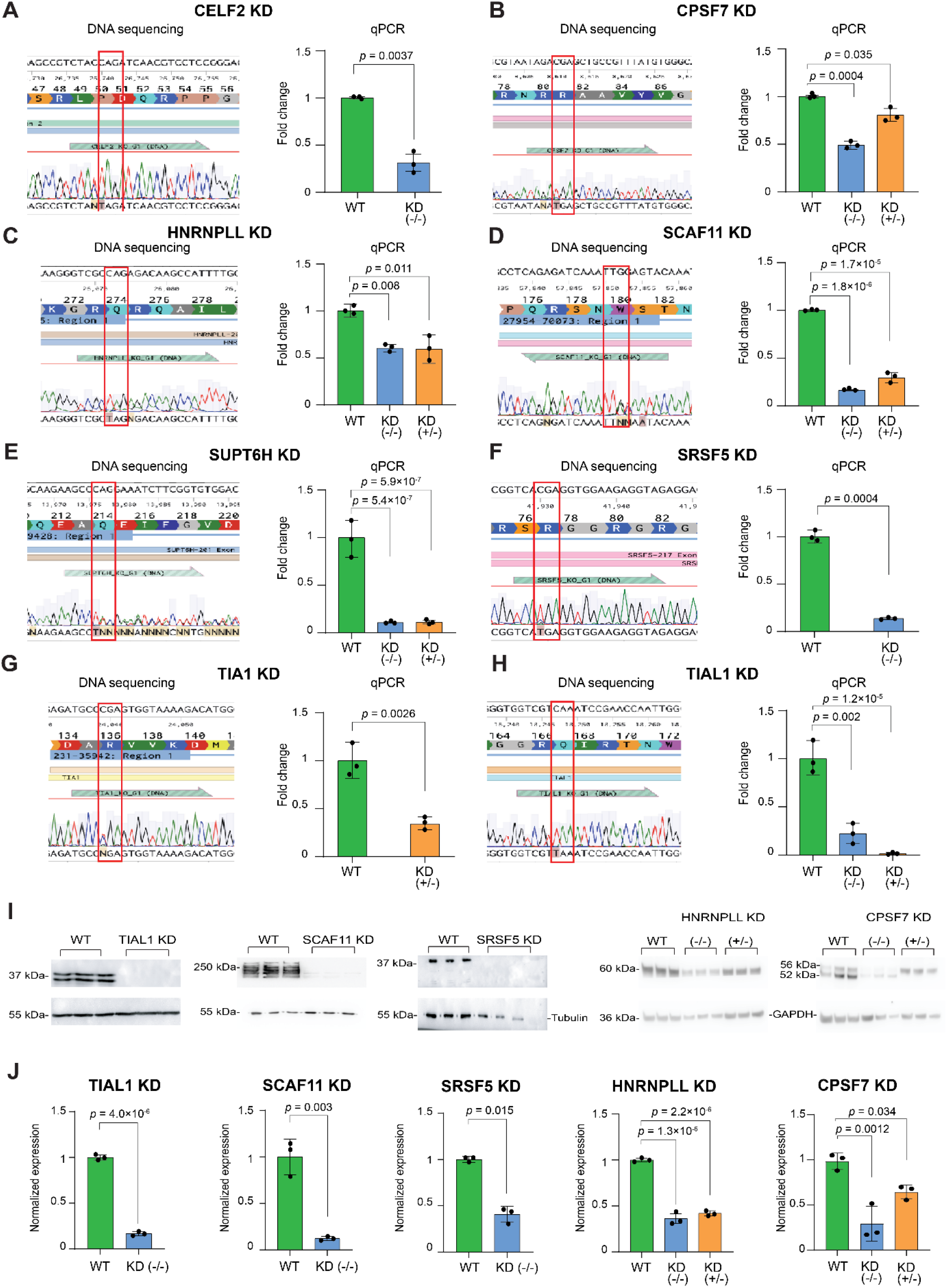
Validation of RBP knockdown by DNA sequencing, qPCR and Western Blot. DNA sequencing results demonstrating C to T edits (red box) resulting in new stop codons TAG, TAA or TGA, and relative RNA fold change (FC) in **(A)** CELF2, **(B)** CPSF7, **(C)** HNRNPLL, **(D)** SCAF11, **(E)** SUPT6H, **(F)** SRSF5, **(G)** TIA1 and **(H)** TIAL1 knockdown (KD) iPSCs compared to control (WT). Homozygous (-/-) edits are shown except for TIA1, where only a heterozygous (+/-) edit was achieved. **(I)** Western blot validation of knockdown efficiency for TIAL1, SCAF11, SRSF5, HNRNPLL and CPSF7 in knockdown (KD) cells compared to WT controls. Tubulin (TIAL1, SCAF11, SRSF5) and GAPDH (HNRNPLL, CPSF7) were used as loading controls. **(J)** Densitometric quantification of protein knockdown levels normalized to their respective loading controls. CPSF7 measurement includes both bands of its characteristic 52/56 kDa doublet (uncropped blots in Supplementary Figure 1b). For qPCR (A-H) and western blots (I, J), *n* = 3 independent experiments per line. Data are presented as mean ± s.d. Each KD line was compared with WT by two-sided Welch’s t-test (two groups) or by one-way ANOVA with two-sided Dunnett’s multiple-comparisons test (three groups). Exact P values are shown in the figure.

**Extended Data Figure 6:**
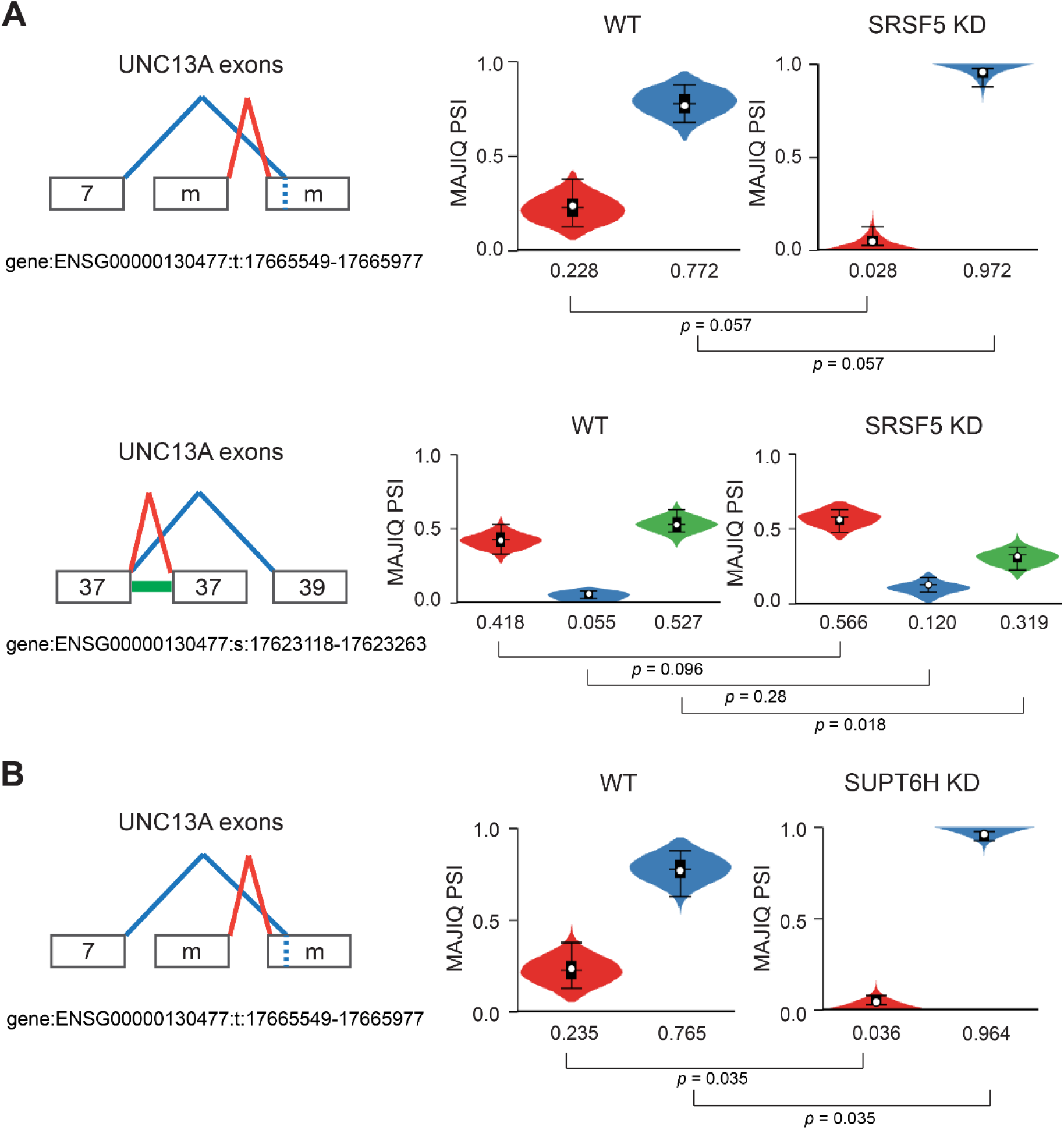
Aberrant splicing of UNC13A upon SRSF5 and SUPT6H knockdown. Output of MAJIQ ΔPSI analysis comparing WT and **(A)** SRSF5 or **(B)** SUPT6H KD neurons, displaying the two events in UNC13A that were significantly altered. Numbers indicate canonically annotated exons, and ‘m’ represents exons identified by MAJIQ. Violin plots show the MAJIQ posterior distribution of PSI for each group (*n* = 4 independent experiments). Boxes represent the 25th-75th percentiles of the posterior distribution, with the median shown as a line; whiskers extend to the 10th and 90th percentiles. The white circles indicate the expected PSI. MAJIQ P values were calculated as 1 − the posterior probability of change (two-sided, no adjustment for multiple comparisons). Exact P values are shown in the figure.

**Extended Data Figure 7:**
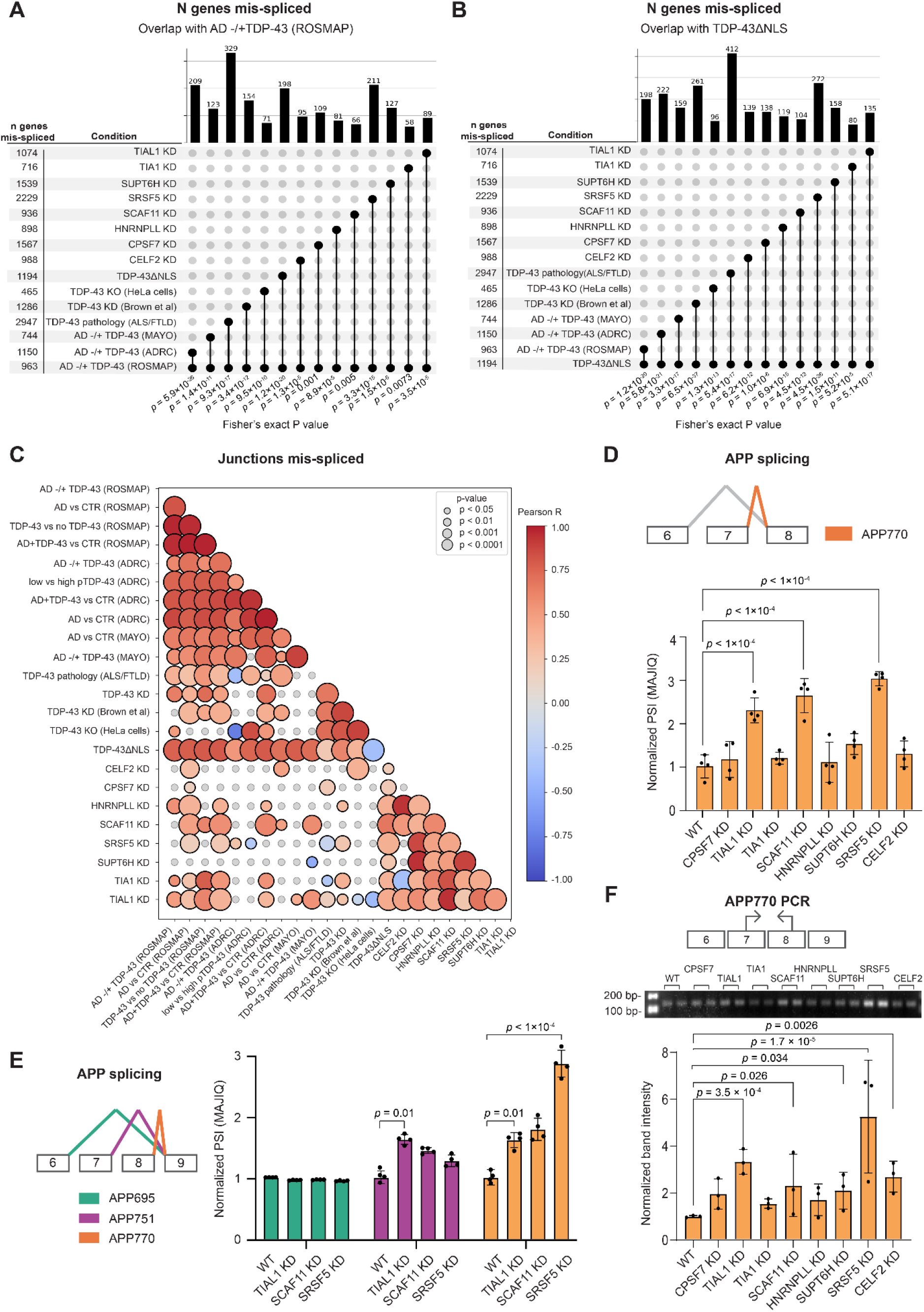
Overlap and concordance of splicing alterations across patient and cell models. Upset plot comparison of shared splicing changes, in patient and cell models to **(A)** TDP-43 pathology in AD in the ROSMAP cohort, and **(B)** TDP-43ΔNLS. Numbers on the left indicate the total number of mRNAs aberrantly spliced, and numbers on the top indicate the number of mis-spliced mRNAs shared between datasets. For cell models, a significance threshold of |ΔPSI| > 0.1 and P-value < 0.1 was maintained; for patient models, a threshold of |ΔPSI| > 0.02 and P-value < 0.05. One-sided Fisher’s exact test (enrichment), no adjustment for multiple comparisons. **(C)** Heat map representing the Pearson correlations of ΔPSI values of splice junction-level events shared between each pair of conditions. Bubble size indicates p-value and hue reflect the Pearson *r* (two-sided test). Grey: non-significant correlations. Aberrant splicing of APP mRNA in induced neurons following KD of the indicated RBPs compared to control. **(D)** MAJIQ PSI values of the APP exon 7-8 junction, producing the APP770 isoform. **(E)** MAJIQ PSI values of APP exons 6-9, normalized to control per junction. In D and E, *n* = 4 independent experiments per line, PSI values were extracted from MAJIQ VOILA; conditions were compared with MAJIQ ΔPSI (*P* = 1 − posterior probability of |ΔΨ| > 0.02), not adjusted for multiple comparisons; **** indicates a MAJIQ *P* < 1 × 10⁻⁴, the smallest value MAJIQ reports. **(F)** Gel densitometry of the APP PCR product, normalized to the WT (KOLF2.1J) control band (n = 3 independent experiments). Size markers (bp) are indicated; the APP770 product is 136 bp (uncropped gel in Supplementary Figure 1c). One-way ANOVA with two-sided Dunnett’s multiple-comparisons test vs WT. Data are presented as mean ± s.d. Exact P values are shown in the figure.

**Extended Data Figure 8:**
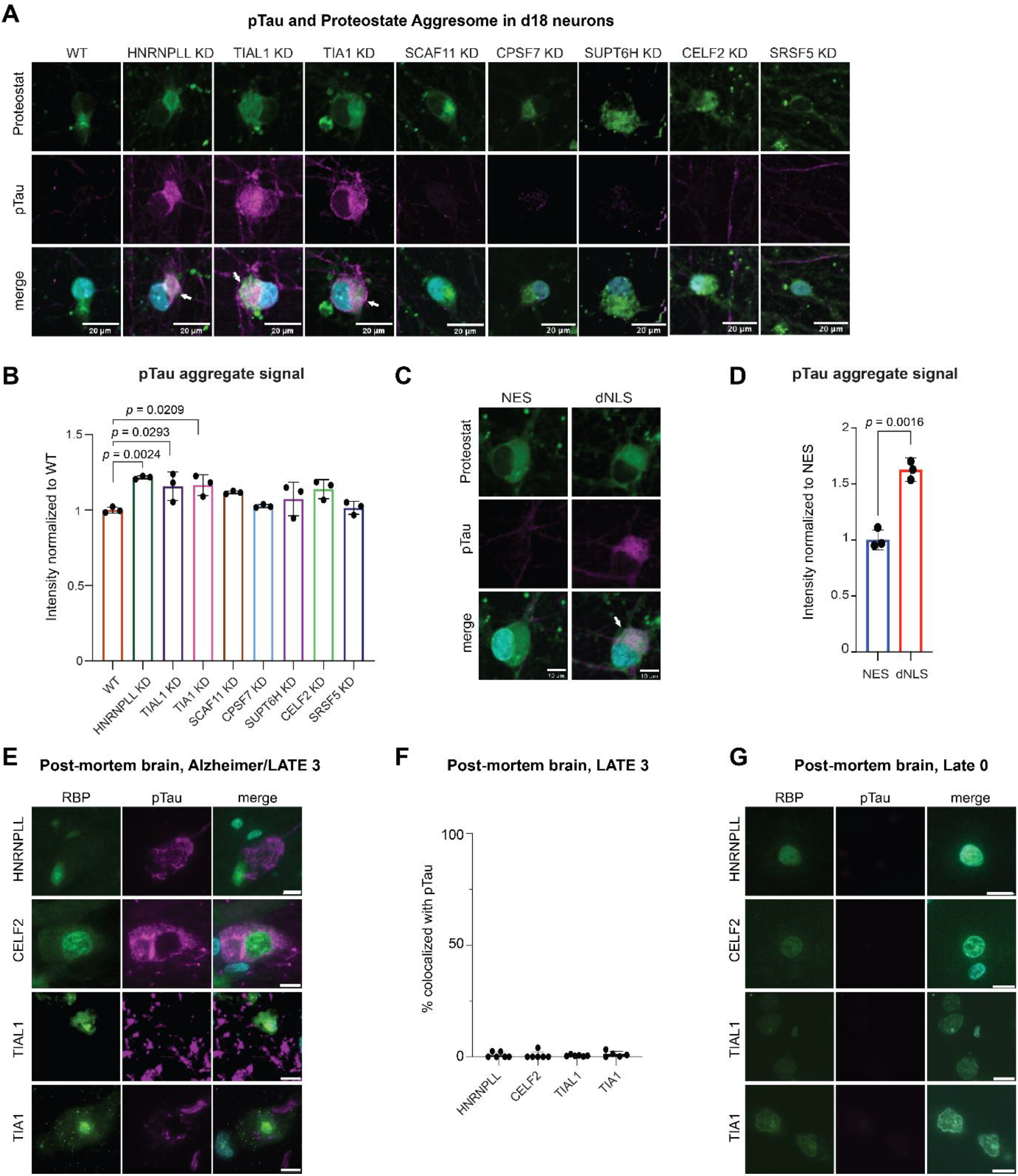
pTau pathology in TDP-43 models and AD/LATE brain(A) Immunofluorescence staining of day 18 (d18) induced neurons following knockdown of target RBPs. Cells were stained for pTau (purple), PROTEOSTAT aggresome detection dye (green) and Hoechst (cyan, nuclei in the merge channel). **(B)** Quantification of pTau mean intensity in aggregates segmented by the PROTEOSTAT aggresome detection dye, normalized to WT (*n* = 3 independent differentiations (dots), each the mean of three technical replicates with 98 images per repeat). Data are presented as mean ± s.d. One-way ANOVA with two-sided Dunnett’s multiple-comparisons test versus WT. Similarly, **(C)** immunofluorescence staining and **(D)** quantification of pTau mean intensity in induced neurons expressing NES (control) or TDP-43ΔNLS, normalized to NES (*n* = 3 independent differentiations). Data are presented as mean ± s.d. Two-sided Welch’s t-test. pTau and RBP of interest co-immunohistofluorescence in **(E)** Alzheimer/LATE3 neuropathological specimens (scale bars: 10µm), and **(F)** percentage of pTau inclusions co-localizing with specified RBPs, per case (mean ± s.d.; each dot represents one case). FFPE sections analyzed for HNRNPLL (*n* = 6), CELF2/CUGBP2 (*n* = 6), TIA1 (*n* = 6) and TIAL1 (*n* = 3). TIAL1 was also evaluated in cryosectioned frozen tissue (*n* = 3). **(G)** Similar immunohistofluorescence on non-neurological disease, control postmortem brains (images representative of 4 control cases per RBP). Exact P values are shown in the figure.

**Extended Data Figure 9:**
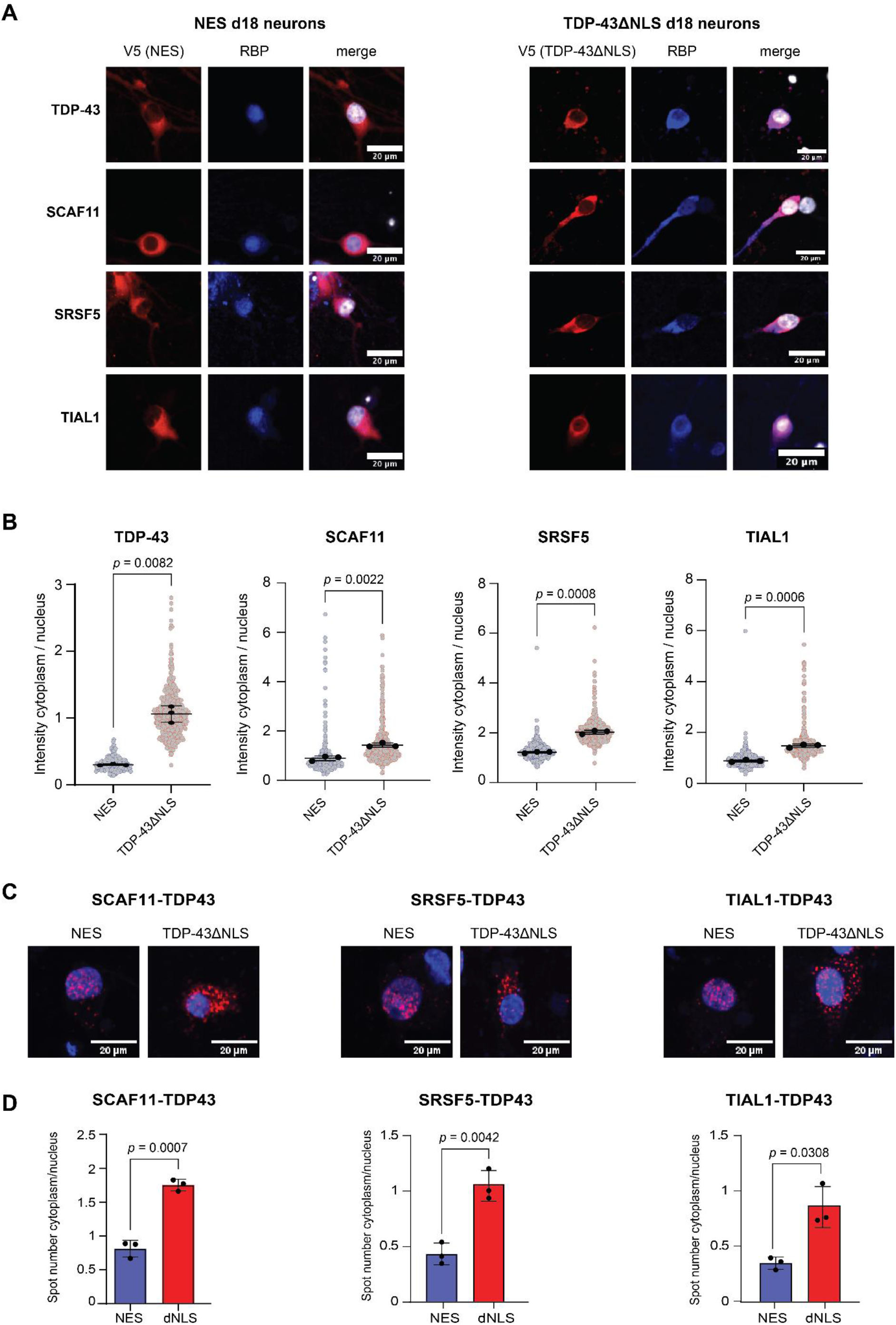
Cytoplasmic mislocalization of TDP-43 promotes aberrant cytoplasmic accumulation of RNA-binding proteins. **(A)** Immunofluorescence staining of day 18 (d18) iPSC-derived neurons expressing NES (control) or TDP-43ΔNLS. Cells were stained for V5 (red), an indicated RNA-binding protein (RBP; blue: TDP-43, SCAF11, SRSF5, TIAL1) and Hoechst (grey, nuclei in the merge channel). Scale bars, 20 µm. **(B)** Quantification of the indicated RBP cytoplasmic-to-nuclear intensity ratio. Black dots represent the mean of each differentiation on which statistics were performed (n = 3 independent differentiations); light dots represent individual cells. Data are presented as mean ± s.d. Two-sided Welch’s t-test, no adjustment for multiple comparisons. **(C)** Representative fluorescence images of nanoscale (<40 nm) TDP-43-RBP proximity (red), by proximity ligation assay (PLA) in control (NES) versus TDP-43ΔNLS models. Hoechst (blue, nuclei). Scale bars, 20 µm. **(D)** Quantification of the cytoplasmic-to-nuclear PLA spot abundance ratio. Data are presented as mean ± s.d.; n = 3 independent differentiations per condition. Two-sided Welch’s t-test, no adjustment for multiple comparisons. Exact P values are shown on the panels.

**Extended Data Table 1:** Splice junctions significantly altered upon RBP KD in iPSC-derived neurons. Significant junctions: |ΔPSI| > 0.1 and P < 0.1 (MAJIQ ΔPSI; P = 1 − posterior probability of change, two-sided; no adjustment for multiple comparisons), comparing each KD line with WT KOLF2.1J neurons (*n* = 4 independent experiments per line). Exact values for each junction are provided in Supplementary Table 2.

| **KD condition** | **Number of junctions that were included more / less upon KD** |
|---|---|
| **SRSF5 (homozygous)** | **2656 /2708** |
| **CPSF7 (homozygous)** | **1666 /1574** |
| **SUPT6H (homozygous)** | **1533 /1559** |
| **TIAL1 (homozygous)** | **1108 /1120** |
| **CELF2 (homozygous)** | **950 /943** |
| **HNRNPLL (homozygous)** | **903 /923** |
| **SCAF11 (homozygous)** | **870 /934** |
| **TIA1 (heterozygous)** | **691 /645** |

## Materials and methods

### Datamining RNA-seq from human cortices

For analysis of AD and control brain tissues with and without TDP-43 pathology, we included bulk RNA-seq data from dorsolateral prefrontal cortex (DLPFC) samples from the Religious Orders Study and Rush Memory and Aging Project (ROSMAP) ^20^ available in Synapse (syn23446022). All participants in ROSMAP were enrolled without known dementia and agreed to annual clinical evaluation and brain donation. Both studies were approved by the Institutional Review Board of Rush University Medical Center. Each participant signed informed and repository consents and an Anatomic Gift Act. Raw bam files from 450 samples (demographics and projids in **Supplementary Table 1**) were used as input for splicing analysis described below. Normalized isoform-level expression data as determined by RSEM ^75^ from 505 samples (**Supplementary Table 1**) was used as orthogonal verification. When comparing samples without TDP-43 pathology to those with TDP-43 pathology, TDP-43 stage (tdp_st4) was dichotomized as per RADC recommendation: no pathology = no TDP-43 pathology or TDP-43 pathology in amygdala only (Stages 0 and 1); TDP-43 pathology = TDP-43 pathology extending beyond amygdala (Stages 2 and 3) ^76,77^. For measurements of Aβ pathology by immunohistochemistry, we used the RADC amylsqrt_est_8reg variable, reflecting mean amyloid score across 8 brain regions ^73,78^ in ROSMAP, excluding samples without any entry for this variable. For correlation of Aβ pathology to *APP* isoforms, we used min-max normalization (0–1 scale) to analyze individual (APP695, APP751, APP770) and combined pathogenic (APP751+770) expression relative to the canonical APP695 isoform. RNA-seq data from *C9orf72* ALS/FTLD-TDP mid-frontal neocortex were obtained from Liu et al. (2019)^29^. Briefly, neuronal nuclei were isolated and FACS-sorted to separate TDP-43-negative (pathological) from TDP-43-positive fractions, prior to RNA extraction and sequencing.

### Datamining proteomics from human cortices and hippocampi

Proteomic data of ROSMAP DLPFC tissues from 700 AD and control subjects (**Supplementary Table 1**) were from ^28,71,79^. ADRC hippocampal proteomics data (90 subjects) were obtained from ^71^. Mass spectrometry and database search followed ^28^, however, both peptide level and protein level searches were retained. Samples from a total of 1124 individuals were normalized using TAMPOR ^80^. Batch effects were removed using a parallelized, ordinary, nonparametric, bootstrapping regression. Only APP peptides were considered, subjected to a 50% missingness filter and checked for outliers. One-way ANOVA was conducted for all APP peptides across TDP-43 level as defined by ROSMAP.

### RNA-seq from human hippocampi and amygdalae

Postmortem hippocampi of 91 cases from the University of Kentucky AD Research Center autopsy cohort were analyzed. Details of the cohort, pathological and clinical diagnosis were described previously ^23–25^. In ref. ^23^, cases with hippocampal pTDP-43 are classified as ‘LATE-NC + ADNC’ and those without as ‘ADNC’ or ‘AD’. RNA was extracted and strand-specific poly(A) selection was performed via oligo(dT) beads prior to reverse-transcription. After quality control (Qubit, PCR, Bioanalyzer), libraries were sequenced on an Illumina NovaSeq 6000 (PE150) at Novogene. Reads were trimmed and aligned to GRCh38 with STAR (v2.7.3a) as described below (RNA sequencing and genome alignment). The Mayo Clinic cohort consisted of 85 amygdala samples ^81^; demographic information is in **Supplementary Table 1**. Bulk RNA-seq reads shorter than 30bp were discarded using *fastp* ^82^. The trimmed FASTQs were aligned to GRCh38 using STAR (v2.7.3a) ^83^ (with EndToEnd option and outFilterMismatchNoverLmax set to 0.04) with gene annotations from Ensembl (release 99) ^84^. For both cohorts, aligned BAMs were used as input for differential splicing analysis with MAJIQ Heterogen (v2.5.7) ^21^. Prior to running MAJIQ Heterogen, MOCCASIN (v0.24) ^85^ was used to adjust MAJIQ build files by RNA integrity number, sequencing run, and flow cell lane.

### Generation of knockdown iPSC lines

KOLF2.1J iPSCs ^86^ were genetically edited using a BE3 cytidine base editor^53^ with minor adjustments to established protocols^87^. Guide RNAs designed via Benchling were cloned into the pKLV2 gRNA plasmid (Addgene #67974) via Gibson assembly. Top off-target sites were sequenced to verify editing specificity (primers in **Supplementary Table 7**). For transfection, 0.5 million iPSCs were seeded in a 3.5 cm plate. The following day, iBE3 CLYBL (Addgene #174569) and guide plasmids (1 µg each) were introduced via TransfeX in mTeSR1 with 10 µM ROCK inhibitor (ROCKi). Post-transfection, cells underwent two days of selection with puromycin (0.8 µg/ml) and doxycycline (2µg/ml). On day three, iPSCs were single-cell seeded into 96-well plates via serial dilution and expanded for ∼2 weeks (with ROCKi for the first 2-3 days). Single cell clones were transferred to 24-well plates, harvested for DNA sequencing, and validated clones were expanded. Finally, the PG-TO-hNGN2 plasmid was integrated as previously described.

### Lentiviral CRISPRi Knockdown of TDP-43 in iPSC-derived neurons

Vector construction and virus production dual-guide constructs targeting the transcription start site of *TARDBP* (TDP-43, CAGGGACACCGAAGCAGCGA; CCTGTCGGGCTTCCCAGCAG) and non-targeting (NT, GGACTAAGCGCAAGCACCTA; GCGGTGGGCTCCCGGACCGC) were synthesized (Twist Bioscience) and cloned into CSM entry vectors via Golden Gate assembly^88^. Following bacterial transformation and plasmid preparation, lentivirus was produced by transfecting 6 µg transfer plasmid, 6 µg psPAX2 (Addgene #12260), and 4 µg MD2.G (Addgene #12259) to 5 × 10⁶ Lenti-X cells (Takara Bio, #632180) in 10 cm dishes, using Lipofectamine 3000. Supernatants collected at 48 h and 72 h were pooled, clarified (500 × g), and concentrated 10-fold using Lenti-X Concentrator (Takara Bio, #631231) before storage at −80°C. CRISPRi Knockdown and Differentiation CRISPR interference (CRISPRi) was performed in iW11 iPSCs stably expressing TO-NGN2 and dCas9-ZIM3. Cells were transduced with CSM-dual-sgRNA lentivirus targeting the *TARDBP* transcription start site. Following transduction, iPSCs were differentiated into cortical-like i3Neurons. Day 18 neurons were harvested for RNA extraction to evaluate knockdown efficiency and perform sequencing.

### Differentiation of iPSCs to neurons

Human iPSCs were cultured and differentiated as described previously ^32,74,89^. Briefly, iPSCs (70-80% confluent) were transferred to Matrigel-coated plates in induction medium (DMEM/F12, N2, NEAA, L-glutamine) with 10μM ROCKi and 2μg/ml freshly prepared doxycycline (DOX). Following two days of DOX induction and one day of DOX + 40μM BrdU (mitotic depletion), on day 3, neurons were dissociated (Accutase) and seeded onto PDL-coated plates. Maturation medium consisted of Neurobasal Plus/B27 Plus supplemented with CultureONE, BDNF/GDNF/NT3 (10 ng/ml), L-ascorbic acid, and laminin. Half-medium changes (minus DOX/ROCKi) were performed on days 7, 10, and 14.

### Inducible TDP-43^ΔNLS^ and NES iPSC-derived neurons

Cytoplasmic TDP-43 mislocalization was modeled with a monoclonal NGN2-iPSC line carrying a doxycycline-inducible TDP-43 construct with a mutagenized nuclear localization signal (NLS) - rendering it cytoplasmic, as described in ref. ^32^. As a cytoplasmic overexpression control, we used neurons carrying a doxycycline-inducible nuclear export signal (NES). Both TDP-43^ΔNLS^ and NES were fused to APEX2 ^90^ to enable proximity labeling. Differentiations into day 8 neurons were achieved with estimated efficiency of > 90%, as confirmed by MAP2 immuno-staining. TDP-43^ΔNLS^ and NES constructs were induced on the last day of differentiation. Expression of TDP-43^ΔNLS^ was minimal without doxycycline. Upon doxycycline induction (2 µg/ml, for 24 hours), cytoplasmic TDP-43 increased by ∼10-fold relative to nuclear TDP-43. Total TDP-43^ΔNLS^ levels were ∼3-fold higher than endogenous TDP-43 in the uninduced condition ^32^. Endogenous TDP-43 levels were ∼2.7-fold higher upon TDP-43^ΔNLS^ induction.

### RNA sequencing and genome alignment

RNA-seq libraries were prepared and sequenced at the Crown Genomics institute of the Nancy and Stephen Grand Israel National Center for Personalized Medicine, Weizmann Institute of Science using the TruSeq protocol (Illumina). RNA was extracted using the miRNeasy Kit (Qiagen) and its concentration quantified with a Nanodrop spectrophotometer. Poly-(A) mRNA fraction was purified from 500 ng of total input RNA, followed by fragmentation and the generation of double-stranded cDNA, Agencourt Ampure XP beads cleanup (Beckman Coulter), A base addition, adapter ligation and PCR amplification. Libraries were quantified by Qubit (Thermo Fisher Scientific) and TapeStation (Agilent). Sequencing was done on Novaseq 6000 S1 300 cycles kit or Novaseq X 1.5B 300 cycles kit, allocating ∼50-60M reads per sample (Illumina; paired end sequencing). In the case of TDP-43^ΔNLS^ and NES samples, sequencing was done with NextSeq high output kit, 75 cycles, allocating ∼20M reads per sample (Illumina; single-end sequencing). Poly-A/T stretches and Illumina adapters were trimmed from RNA sequencing reads using cutadapt ^91^; resulting reads shorter than 30bp were discarded. RNA-seq generated in this study or from published datasets ^10,29,31^, were mapped to the H. sapiens reference genome GRCh38 using STAR ^83^, with Ensembl gene annotations (with EndToEnd option and outFilterMismatchNoverLmax was set to 0.04). Since ROSMAP bam files were aligned to hg19, they were first converted back to *fastq* files before aligning to GRCh38. Reads were grouped based on unique molecular identifiers (UMI), using PICARD MarkDuplicate with BARCODE_TAG parameter at snakemake ^92^.

### Differential gene expression and splicing analysis

Differential gene expression analysis was done with the User-friendly Transcriptome Analysis Pipeline (UTAP) ^93^, whereby normalization of the counts and differential expression analysis was performed using DESeq2 ^94^ (v1.36.0, betaPrior=True, cooksCutoff=FALSE, independentFiltering=FALSE). Raw P values were adjusted for multiple testing using the procedure of Benjamini and Hochberg ^95^. Splicing analysis was performed using MAJIQ (v2.5.7) ^21^. Biases in analysis of human postmortem datasets, stemming from RNA quality (RIN), sequencing batch, platform, and preparation method were corrected using MOCCASIN (v0.24) ^85^. Previously-reported TDP-43-dependent junctions were from ^20,29,31,62,63^, including STMN2 ^9–12^, *MYO18A* ^62^ and *ANXA11* ^63^. To identify significant splicing variations between groups, we applied a ΔPSI threshold of 2% and a posterior probability of change > 0.9 for the analysis of cell lines or isolated human brain nuclei ^29^ using the MAJIQ ΔPSI function. In post-hoc analyses, a ΔPSI threshold of 10% and P-value threshold of 0.1 was used to determine significant events. Nominal P-values were defined as ‘1’ minus the posterior probability of a significant splicing change. To account for batch effects across RNA-seq runs, RNAseq of RBP KD lines (Extended Data Figure 7D-E) were visualized by normalizing MAJIQ PSI values to their respective batch-matched controls. Human postmortem datasets were compared using MAJIQ Heterogen, which treats each sample as an independent replicate and applies two-sample tests to per-sample PSI posterior distributions, applying a ΔPSI threshold of 2% and a P-value threshold of 0.05. Significance was assessed with the two-sided TNOM test, with P values reported as the median across posterior samples, without adjustment for multiple comparisons. Post-hoc significant events were called using the same thresholds, where ΔPSI was calculated as the difference in median PSI between disease and control cohorts. Voila TSV output tables were parsed to extract per-junction values rather than junction clusters, using a modified R script based on previously described methods ^10^ (see Code Availability). To identify the significance of overlap in mis-spliced genes between groups, we used one-sided Fisher’s exact test. Significance of three-way overlap was assessed using a permutation test, in which gene sets of equal size were randomly sampled from the common background (10,000 permutations). The empirical p-value was calculated as the fraction of permutations yielding a triple overlap greater than or equal to the observed overlap. For visualization of individual splice events, the ‘MANE select’ transcript ^96^ was used for annotation of exon numbers. PSI values were calculated per splice junction block as defined by MAJIQ, summing to 1 within each block. Junctions were matched across datasets by unique MAJIQ local splicing variation identifiers.

### Spatial transcriptomics

Anterior hippocampal sections from 12 cases with LATE-NC stage 0 (no TDP-43 pathology), and 12 with LATE-NC stage 2 or 3 were analyzed (demographics in Supplementary Table 1). Xenium in situ spatial transcriptomics was performed using a 480-plex probe panel, including probes targeting cryptic exons in 77 transcripts. Cells were included if they fell within the sample-specific 1st-99th percentile range for cell area, had a nucleus-to-cell area ratio between 0.02 and 0.95 and contained at least two detected mRNAs. No minimum total-transcript threshold was imposed. mRNAs detected in fewer than three cells were excluded. Not all cell types were detected in every donor; donors lacking a given cell type were excluded from the corresponding analyses (per-panel *n* values are given in the figure legends). Neurons were defined by the expression of either NeuN/RBFOX3 or of at least two neuronal markers (*SNAP25, NRGN, SLC17A7, GAD1*) and by the morphology-based criteria ^97^ (probability of ≥ 0.80, ≥ 99.8% specificity). Neuron type identities were assigned using canonical marker-gene expression: dentate granule cells, *RBFOX3+, SLC17A7+, PROX1+, CALB1+, GAD1/2-*; pyramidal neurons, *RBFOX3+, SLC17A7+, PROX1-, ZBTB20+*; mossy cells, *RBFOX3+, SLC17A7+, CALB2+, PROX1-, GAD1/2-*; basket cells, *RBFOX3+, GAD1/2+, CALB1+, TAC1+, PVALB+, SLC17A7-*; chandelier cells, *RBFOX3+, GAD1/2+, TAC1-, SST-, SLC17A7-*; O-LM cells, *RBFOX3+, GAD1/2+, SST+, LHX6+, SLC17A7-*; and Cajal-Retzius cells, *RBFOX3+, SLC17A6+, SLC17A7-, GAD1-, GAD2-*. Neurons with at least one *STMN2* cryptic-exon transcript ^11,12^ were classified as *STMN2* CE⁺, given the sparse detection of cryptic-exon probes. IHC sections were manually registered to the matched Xenium DAPI image using a rigid transform. TDP-43+ cells were detected in the IHC using color thresholding in HSV space (hue 0.38– 0.55, saturation ≥0.28, value ≤0.92, area 150–15,000 px²). Burden was defined as TDP-43+ cells per mm² of tissue, derived from the DAPI mask. For the spatial analysis, *APP* transcripts were binned onto a 50×50 µm grid covering the full section, generating separate density heatmaps for the long isoform (exon 7/8) and short isoform (exon 6-9 junction), from which a local ratio heatmap was computed as 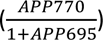. TDP-43+ cell centroids were binned onto the same grid to produce a TDP-43 density heatmap, and Spearman’s ρ between the *APP* ratio and TDP-43 density was computed per donor across all bins with at least one transcript. LATE stages 2 and 3 were pooled and compared to LATE 0 using a two-sided Mann-Whitney U test. To compare long/short *APP* isoform ratio values between groups, we used linear mixed-effect modeling, with Satterthwaite’s formula for degrees of freedom estimation, via lme4 and lmerTest in R ^98–100^. The grouping variable (e.g., LATE stage or *STMN2* cryptic exon status) was included as a fixed effect, and donor was included as a random intercept to account for the non-independence of multiple cells measured from the same subject. Models were fitted using maximum likelihood estimation. For cell-level analyses, the per-cell ratio was calculated as APP751/770 counts divided by APP695 counts in cells with at least one APP695 transcript; per-donor values were the ratio of summed APP751/770 to summed APP695 counts.

### Subcellular fractionation

Cells lysed with buffer A (10 mM Tris Cl, pH 7.5; 10 mM NaCl; 3 mM MgCl2; 0.5% NP40) with cOmplete Protease Inhibitor Cocktail (Roche, 4693116001) and PhosSTOP (Roche, 4906837001), incubated on ice for 10 min with intermittent vortexing and centrifuged to pellet nuclei (10 min at 4**°**C). The cytoplasmic supernatant was collected. Nuclei were washed repeatedly with buffer A until resuspension ceased. Pellets were then resuspended in RIPA buffer (150mM NaCl, 1% NP40, 0.5% DOC, 0.1% SDS, 50mM Tris pH 8), sonicated (3 cycles, 10s on/20s off, 30-40% power), and centrifuged to obtain the nuclear fraction. Nuclear and cytoplasmic fractions underwent in-solution tryptic digestion via the S-Trap method ^101^. Briefly, lysates were heated (96°C, 5 min) and sonicated (6 cycles, 30s, (Bioruptor Pico, Diagenode)). After BCA protein quantification (ThermoScientific), 20 µg of protein per sample were reduced (5 mM DTT) and alkylated (10 mM iodoacetamide). Samples were loaded onto S-Trap microcolumns (Protifi), washed (90% methanol/50 mM ammonium bicarbonate), and digested with trypsin (1:50) for 1.5h at 47°C. Following ammonium bicarbonate elution, additional trypsin (1:50) was added for overnight incubation at 37°C. Peptides were further eluted with 0.2% formic acid (in H2O and 50% acetonitrile), pooled, and vacuum-dried. Samples were desalted via solid-phase extraction (Oasis HLB, Waters), vacuum-dried, and stored at −80°C.

### Liquid chromatography, mass spectrometry and proteomics

Liquid chromatography and mass spectrometry and proteomic analysis were performed as described in ^102^. Peptides were separated via nanoUPLC (nanoAcquity, Waters) using a C18 trap and a T3 HSS nano-column (250 mm). A 105-min gradient (4–30% B; 0.35 µL/min) eluted peptides into an Orbitrap Exploris 480 (Thermo Scientific) via a nanoESI emitter. Data were acquired in DDA mode (2s cycle). MS1 resolution was 120,000 (380–1500 m/z); MS2 resolution was 15,000 with a 1.4 m/z isolation window and 40s dynamic exclusion. Raw data were analyzed via MaxQuant (v1.6.6.0)^103^, using the Andromeda search engine against the UniProt human database. Search parameters included trypsin specificity (up to 2 missed cleavages), fixed carbamidomethylation, and variable methionine oxidation/N-terminal acetylation. False discovery rate (FDR) was set to 1% for peptides and proteins. Label-free quantification (LFQ) and “match-between-runs” were enabled. Downstream analysis in Perseus (v1.6.2.3)^104^ involved log2-transformation and exclusion of contaminants, reverse proteins, and “only identified by site” entries. Data were filtered for at least 6 valid values per row, with remaining missing values imputed from a normal distribution. Two-sample t-test comparing samples with biotin to samples without biotin was performed to remove non-specific binders (FDR < 0.05, log2FC > 0). Samples were separated based on stress condition before filtering to remain only those proteins with a minimum of 2 valid values in at least one experimental group (TDP-43^ΔNLS^ or NES). Missing values were imputed again using a normal distribution. Two-sample t-test using log2-transformed LFQ values was performed to gain proteins enriched in TDP-43^ΔNLS^. Functional enrichment analysis was performed using PANTHER Overrepresentation Test against the Gene Ontology database ^105^ with all experimentally identified proteins serving as the reference list.

### Semi-quantitative PCR and specific transcript splicing

Total RNA was extracted from day 8 differentiated cells using the miRNeasy MicroKit (Qiagen). RNA (500 ng) was reverse-transcribed via the High-Capacity cDNA Kit (Thermo, 4368814). PCR was performed using primers capturing APP770 CCCATTCTTTTACGGCGGATG; AACAGGATCTCGGGCAAGAGG or *APP* exons 6–9 Primers: TCTGTGGAAGAGGTGGTTCG; GGCCTCAAGCCTCTCTTTGG with RedTaq Ready Mix (Sigma, R2523) at 95°C (5 min); 28 cycles of 95°C (30s), annealing (30s), and 72°C (1 min); and a 5 min final extension. Products were resolved on a 1.8% agarose gel, 120V 1 h. Primers spanning exons 6-9 resolve *APP* isoforms based on product size (APP695: 132 bp; APP770: 357 bp; APP751: 300 bp), enabling relative quantification by band intensity.

### Immunofluorescence of cultured neurons

60,000 iPSC-derived neurons per microwell in 96-well plates (Brooks, MGB096-1-2-LG-L), cultured until day 8 or 18, fixed (4% PFA in 1X PBS++, 15 min), washed (PBS), incubated in 0.1% TritonX (10-15 min) and blocked with CasBlock (10 min, RT). Two staining panels were performed. Panel 1, cells were incubated with anti-Aβ, clone 6E10 (BioLegend 803004, 1:500) and Thioflavin S (Sigma-Aldrich T1892) and panel 2, with anti-pTau (Ser202/Thr205), clone AT8 (Invitrogen, MN1020, 1:500). Primary antibodies were diluted in CasBlock and incubated overnight at 4°C, followed by washes (PBS, 2 x 5 min). Cy5-conjugated donkey anti-mouse IgG (Jackson ImmunoResearch, 715-175-151; 1:200) in PBS incubated (1 h, RT) in the dark and washed (PBS) prior to adding Hoechst (ThermoFisher 62249, 1:8000 in PBS) and PROTEOSTAT aggresome detection dye (Enzo ENZ-51035) (20±10 min RT) for panel 2, prior to final washes (PBS). Confocal images were acquired on the Opera Phenix Plus High-Content Imaging System (nuclei (Hoechst), Thioflavin S /PROTEOSTAT aggresome detection dye (Alexa Fluor 488), anti-Aβ/anti-pTau (Cy5)) and analyzed with Harmony PhenoLOGIC software (v5.2). The ‘Find Image Region’ function in Harmony was used to detect aggregate structures within Alexa 488-positive regions, within which anti-Aβ/anti-pTau signal on the Cy5 channel was quantified. Mean anti-Aβ/anti-pTau intensity colocalized within Alexa 488-positive regions was the primary readout, normalized to data from isogenic KOLF controls within each culture batch. Statistical comparisons were performed using One-way ANOVA with two-sided Dunnett’s test (GraphPad Prism v10.6.1), across three independent experiments, each comprising three technical replicates.

### Western Blot

Cells were washed in 1× PBS and lysed in ice-cold RIPA buffer supplemented with cOmplete Protease Inhibitor Cocktail (Roche, 4693116001) and PhosSTOP (Roche, 4906837001). The lysates were cleared by centrifugation at 15,000 g for 10 min at 4 °C. Protein concentrations were quantified using the BCA Protein Assay Kit (Thermo Fisher Scientific, 23227), and 30μg of total protein per well was resolved by SDS–PAGE at 100–120 V for 70 min. Proteins were transferred to nitrocellulose membranes (Whatman, 10401383) at 250 mA for 70 min. Membranes were stained with Ponceau (Sigma, P7170), blocked for 1 h at room temperature with 5% milk in TTBS (Bio-Lab, 002089232300) and incubated overnight at 4 °C with primary antibodies in antibody solution (5% albumin, 0.02% sodium azide and five drops of phenol red in 0.05% TBST): CPSF7 (Proteintech, 55195-1-AP; 1:1,000), HNRNPLL (Proteintech, 26769-1-AP; 1:1,000), SCAF11 (Sigma-Aldrich, HPA045456; 1:200), SRSF5 (Invitrogen, PA5-60383; 1:200), TIAR/TIAL1 (Cell Signaling, 8509; 1:1,000), α-tubulin (Sigma-Aldrich, T9026; 1:4,000) and GAPDH (Invitrogen, AM4300; 1:4,000). Membranes were washed three times for 5 min at room temperature with TBST, incubated for 1 h at room temperature with horseradish peroxidase-conjugated species-specific secondary antibodies, washed three times for 5 min in TBST, and visualized using WESTAR Nova 2.0 (Cyanagen, XLS071) on an ImageQuant LAS 4000 (GE Healthcare Life Sciences). Loading controls were probed on the same membranes as the target proteins. Band intensities were quantified with Fiji (ImageJ v1.54f), normalized to the loading control and expressed relative to the WT mean. Uncropped and unprocessed scans of all blots are provided in Supplementary Figure 1b.

### Proximity Ligation Assay

60,000 iPSC-derived neurons per microwell in 96-well plates (Brooks, MGB096-1-2-LG-L) were cultured until day 18. Cells were fixed (4% PFA in 1× PBS++, 15 min), washed (PBS), permeabilized in 0.1% Triton X-100 (10–15 min) and blocked with CasBlock (10 min, RT). In situ protein–protein interactions were detected using the NaveniFlex Cell kit (Navinci, 60025) according to the manufacturer’s protocol. Wells were incubated overnight at 4 °C with primary antibodies (mouse anti-TDP-43, clone 2E2-D3 (Abnova, H00023435-M01; 1:2,000), paired with rabbit anti-SRSF5 (Invitrogen, PA5-60383; 1:200), anti-TIAL1 (Cell Signaling, 8509; 1:1,000), anti-SCAF11 (Sigma-Aldrich, HPA045456; 1:200), or rabbit anti-TDP-43 (Proteintech, 10782-2-AP; 1:1,000) as a positive control), diluted in Naveni Primary Antibody Diluent. Following PBS-T washes, secondary Navenibodies (anti-mouse M1 and anti-rabbit R2) were applied for 60 min at 37 °C. Ligation, rolling circle amplification and fluorophore hybridization were performed sequentially. Nuclei were stained with Hoechst. Confocal images were acquired on the Opera Phenix Plus High-Content Imaging System, and PLA spots per cell within the nucleus or cytoplasm were segmented and quantified using an image analysis pipeline in Harmony PhenoLOGIC (v5.2).

### Mediation analysis

Mediation analysis was conducted to examine whether *APP* mediates the relationship between TDP-43 pathology and amyloid-β pathology. When modeling different isoforms of *APP* into a single ratio variable 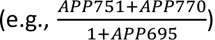, each *APP* isoform underwent min-max scaling before computing the sum/ratio. The total, direct, and indirect effects were estimated using Ordinary Least Squares (OLS) regression models. First, we assessed the effect of TDP-43 on *APP* (path *a*). Next, we examined the effect of *APP* on amyloid-β while controlling for TDP-43 (path *b*). The total effect (*c*) of TDP-43 on amyloid-β was decomposed into the direct effect (*c*′, TDP-43’s effect on amyloid-β independent of *APP*) and the indirect effect (*a* × *b*), representing mediation. To test the statistical significance of the indirect effect, we used percentile bootstrap confidence intervals (95% CI) with n = 1,000 resamples. The proportion of the total effect mediated was computed as 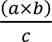. All statistical analyses were performed in Python 3.8.5 using statsmodels (v0.14.1), and results were considered statistically significant at p < 0.05.

### Immunohistofluorescence of human postmortem brain samples

De-identified postmortem brain tissue middle frontal gyrus formalin-fixed, paraffin-embedded (FFPE) tissue blocks were obtained from the Mayo Clinic Florida Brain Bank for Neurodegenerative Disorders with IRB approval and informed consent (Demographics: **Supplementary Table 1**). Neuropathological diagnoses were confirmed by an expert (D.W.D.) using established criteria for FTLD-TDP and LATE ^3,106^. 5 µm sections were deparaffinized in xylene, rehydrated via graded ethanol and subjected to heat-induced epitope retrieval in Citrate (pH 6) or Tris-EDTA (pH 9.0). Sections were blocked/permeabilized (10% NGS, 0.3% Triton X-100 in TBS, 1 h RT) and automated staining was performed using an Opentrons Flex robot with an Omni-Stainer S12 (Parhelia Biosciences). Primary antibodies (**Supplementary Table 8**) were incubated (overnight, 4°C in 5% NGS). After TBS-T washes, secondary antibodies were applied (1 h, RT). Following DAPI counterstaining, slides were treated with 1X TrueBlack (in 70% ethanol) to quench autofluorescence and mounted with ProLong Glass Antifade Mountant (Invitrogen, P36980). Automated imaging was performed on a Nikon Eclipse Ti2 (Spectra X LED; Zyla 4.2 PLUS sCMOS) using NIS-Elements HC. Colocalization of pTDP-43 with RBPs was manually quantified by a blinded investigator. Analysis included all cells within each field across six cases (11– 22 images analyzed per case and 27–64 aggregates quantified per case). Parallel staining of non-neurological disease control autopsies validated the baseline localization of the studied RBPs, ensuring results were not confounded by non-specific staining or experimental artifacts. For TIAL1 immunostaining, frozen frontal tissue punches from *C9orf72* ALS/FTLD-TDP cases were obtained from the Edinburgh Brain Bank (with approval from the Institutional Review Board of the Weizmann Institute of Science (protocol 3046-1); demographics: **Supplementary Table 1**) and cryosectioned (10 μm). Four consecutive sections were mounted per charged glass slide (two disease and two healthy control samples) and stored at −80°C. Slides were fixed overnight at RT in 70% ethanol, 4% formaldehyde and 5% acetic acid, stained for pTDP-43 (22309-1-AP, Proteintech; 1:1000) and TIAL1 (66907-1-Ig, Proteintech; 1:250) using the Opal workflow (Akoya Biosciences). Multi-spectral imaging was performed using the PhenoImager (Opal) system. Protein co-localization was manually assessed in 70 pTDP-43 inclusions per case.

### Statistics and software

Statistical analyses (t-tests, one-way and two-way ANOVA with multiple-comparisons tests) and data plotting were performed with GraphPad Prism (v10.6.1), unless otherwise stated. Fisher’s exact tests, permutation tests, Pearson and Spearman correlations and Mann-Whitney U tests were performed in Python (v3.10.14) using SciPy (v1.14.0), pandas (v2.2.3) and NumPy (v2.2.2); plots were generated with matplotlib (v3.10.0) and ggplot2 (v3.5.1). Linear mixed models were fitted in R (v4.4.1) using lme4 (v1.1-35.5) and lmerTest (v3.1.3), and mediation analyses in Python (v3.8.5) using statsmodels (v0.14.1).

### Data availability

Proximity labeling proteomics data have been deposited to the ProteomeXchange Consortium via the PRIDE ^107^ partner repository with the dataset identifier PXD063021, and fractionation mass spectrometry data are available under PXD063188.

Raw RNA-seq data are available in Sequence Read Archive under accession number PRJNA1255884. Amygdala RNA-seq from the work of Wang et al., ^81^ is deposited to GEO accession GSE324851. Xenium spatial transcriptomics data have been deposited in GEO under accession GSE352490.

ROSMAP ^20^ RNA-seq, proteomics and clinicopathological data (Synapse ID syn23446022) and University of Kentucky ^23^ hippocampal RNA-seq data (Synapse ID syn65658835) are available through the Synapse AD Knowledge Portal (https://www.synapse.org), subject to a data use agreement.

Published datasets reanalyzed in this study are available in GEO under accessions GSE126543 (*C9orf72* ALS/FTLD-TDP neuronal nuclei ^29^) and GSE136366 (TDP-43 knockout HeLa cells ^31^), and in the European Nucleotide Archive under accession PRJEB42763 (TDP-43 knockdown in iPSC-derived neurons ^10^).

Gene Ontology annotations ^33,34,105^ were obtained from release DOI 10.5281/zenodo.7942786, and gene annotations from Ensembl (release 99) ^84^.

## Code availability

Code for running MAJIQ and plotting the results can be found on our GitHub page: https://github.com/WelmoedVZ/splicing.git

## Acknowledgements

We wish to thank Michael Ward (NINDS, NIH) and Bill Skarnes (Jackson labs) for access to iPSC lines. We thank Alon Savidor (WIS) and Yishai Levin (WIS) for mass spectrometry, Sapir Havusha Laufer (WIS) for advice on the fixation protocol for postmortem immunostaining, and Inbal Bolocan Nachman (WIS) and Yuval Elazari (WIS) for RNA sequencing. We also thank Noa Wigoda (WIS), Melina Klostermann (Julius-Maximilians-Universität Würzburg) and the BioCiphers team. Yoseph Barash (UPENN) for bioinformatics advice and the development of MAJIQ. Naomi Habib and Anael Cain (HUJI) for advice and discussions. We thank Anna-Leigh Brown (UCL), Jack Humphrey (MSSM) and Pietro Fratta (UCL) for advice on splicing and generously sharing resources. Emma Marie Monte, Chenchen Zhu (Stanford) for technical assistance with spatial transcriptomics work. EH is the Mondry Family Professorial Chair and Head of the Andrea L. and Lawrence A. Wolfe Family Center for Research on Neuroimmunology and Neuromodulation.

## Author contribution

W.v.Z., T.M. and E.H. conceived and designed the study. W.v.Z., T.M., N.S.Y., E.A., A.R., L.A., E.W., M.N., A.St., R.S.K., M.W., D.M., J.G.M., W.R., M.M.R., A.N.T., Z.T.M. and J.C.-K. analysed the data. W.v.Z., T.M., C.B., A.Si., B.A., L.M., E.A., A.R., D.M., J.G.M. and W.R. performed experiments and molecular analyses. W.v.Z., T.M., L.A., C.S., K.M., A.S.B., Y.W., D.A.B., B.Z., L.J., J.B., M.P.S., M.M.R., D.W.D., M.P., P.T.N., A.N.T., M.C., Z.T.M., N.T.S. and J.C.-K. provided human tissue, samples and cohort resources. W.v.Z., T.M., N.S.Y., A.St., R.S.K., M.W., M.M.R., D.W.D., M.P., P.T.N., J.D.G., A.N.T., M.C., Z.T.M. and N.T.S. generated and curated datasets. W.v.Z., T.M. and E.H. wrote the manuscript that was approved by all authors before submission. E.H. supervised the study and handled correspondence. All authors reviewed and edited the manuscript.

