## Supplementary Figure 1 for "TDP-43 toxic gain of function links ALS, FTD and Alzheimer’s disease via mis-splicing"

### Supplementary Figure 1a: Uncropped gel for Figure 4F

RT-PCR of APP exons 6–9 in NES and TDP-43ΔNLS neurons

L | NES 1-6 | TDP-43ΔNLS 1-6 | L

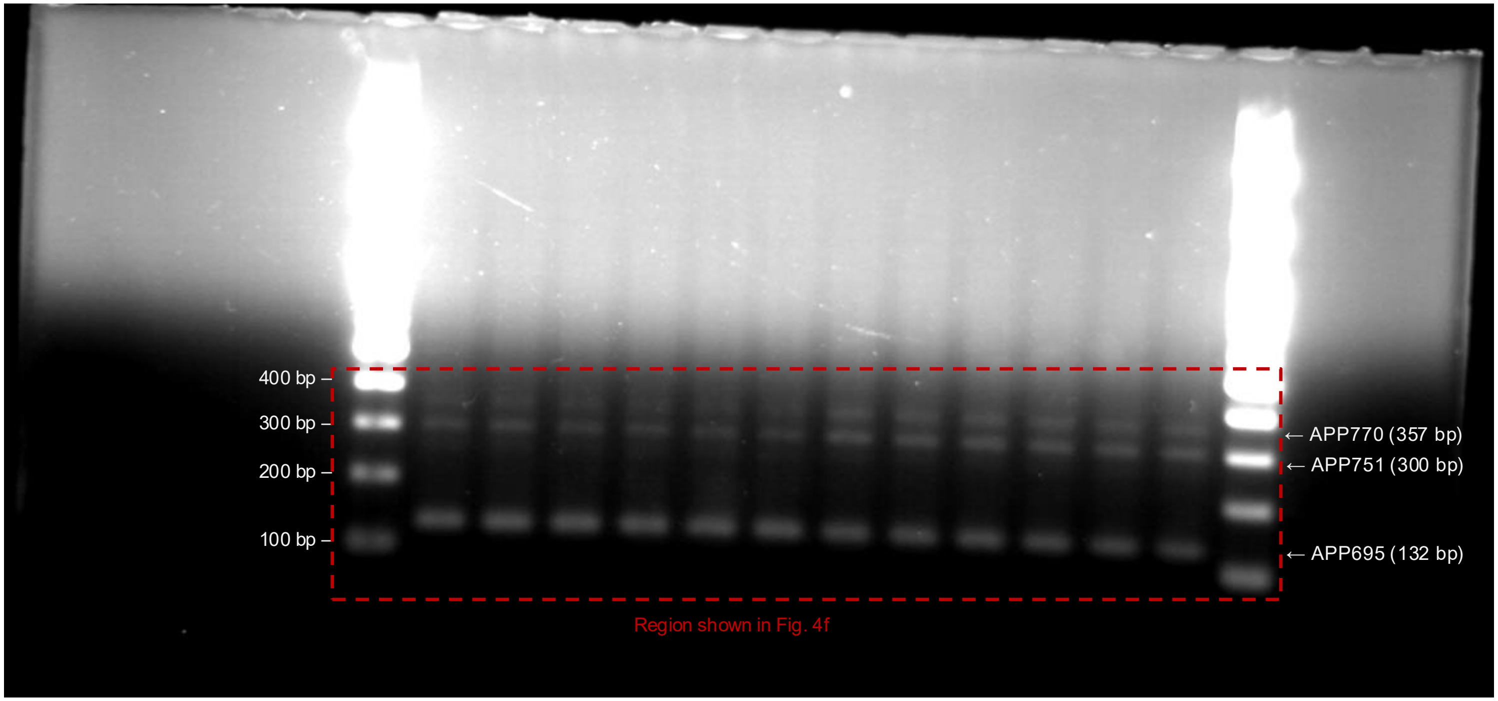


Uncropped image of the RT-PCR gel shown in Figure 4F (primers spanning APP exons 6-9).
L, 100 bp ladder; lanes 1-6, NES; lanes 7-12, TDP-43ΔNLS (n = 6 independent differentiations each). All samples were run on the same gel.
The dashed box indicates the cropped region. Band identities: APP695, 132 bp; APP751, 300 bp; APP770, 357 bp.

### Supplementary Figure 1b: Uncropped blots for Extended Data Figure 5l (1/2)

SCAF11, SRSF5 and TIAL1 knockdown with tubulin loading controls

**SCAF11 KD & WT: SCAF11 antibody**

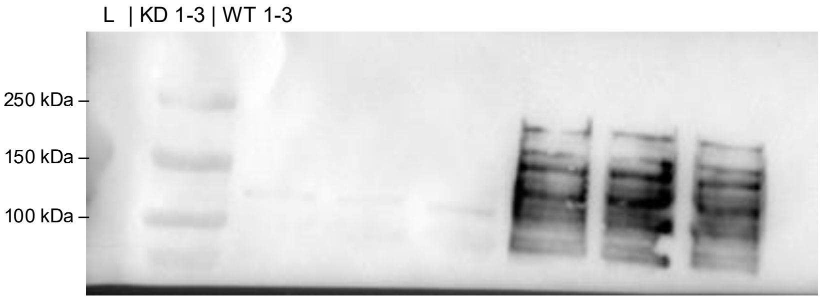


**SCAF11 KD & WT: tubulin (same membrane)**

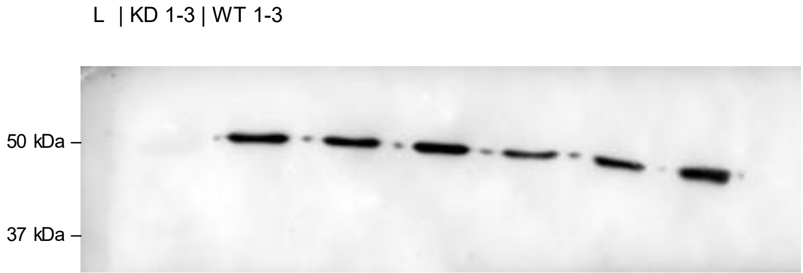


**SRSF5 KD & WT: SRSF5 antibody**

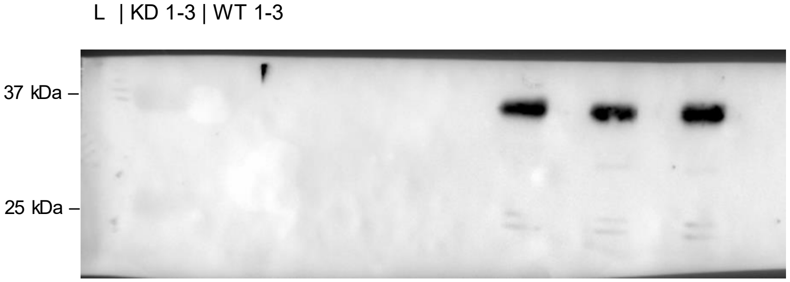


**SRSF5 KD & WT: tubulin (same membrane)**

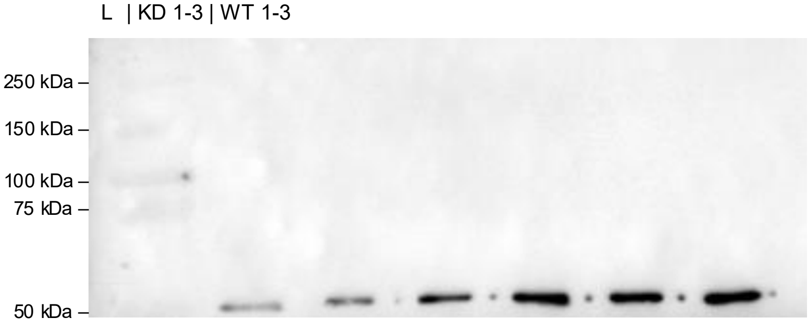


**TIAL1 KD & WT: TIAL1 antibody**

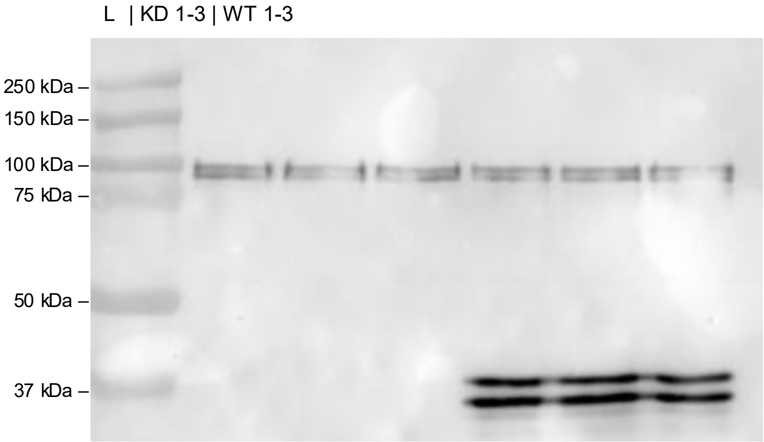


**TIAL1 KD & WT: tubulin (same membrane)**

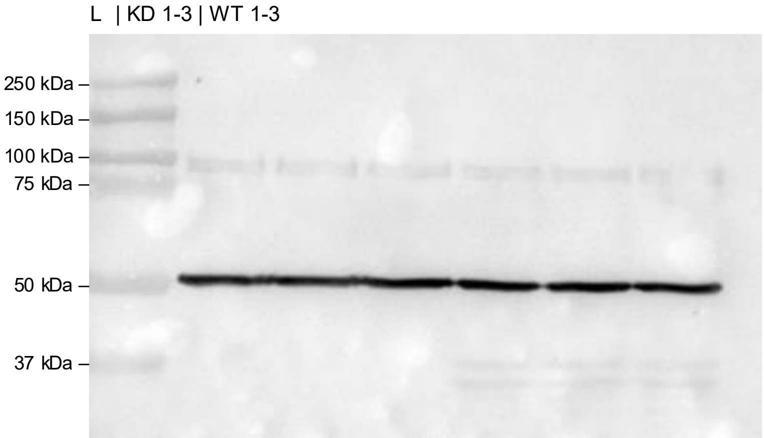


Uncropped western blots for Extended Data Figure 5l. Each membrane was probed for the indicated RBP and then for the loading control (α-tubulin) on the same membrane.
Images of SCAF11, SRSF5 and TIAL1 blots shown in Extended Data Figure 5l were flipped horizontally so that WT appears first.
L, protein ladder (10-250 kDa, Precision Plus Protein Dual Color Standards #1610374); WT and KD, n = 3 independent experiments each.

### Supplementary Figure 1b: Uncropped blots for Extended Data Figure 5I (2/2)

HNRNPLL and CPSF7 knockdown with GAPDH loading controls

**HNRNPLL KD & WT: HNRNPLL antibody**

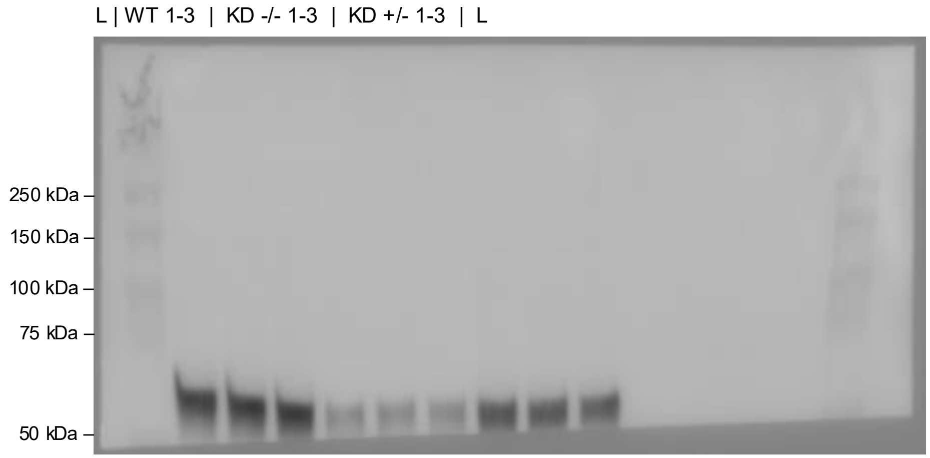


**HNRNPLL KD & WT: GAPDH (same membrane)**

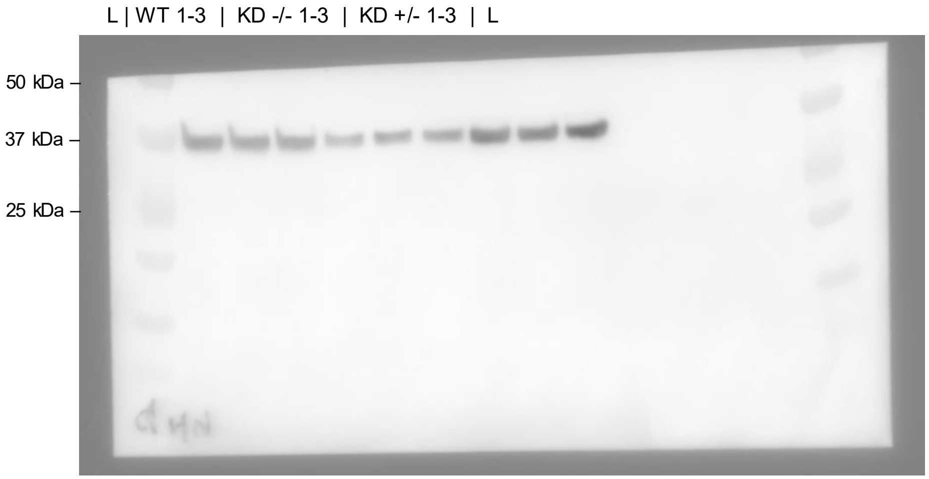


**CPSF7 KD & WT: CPSF7 antibody**

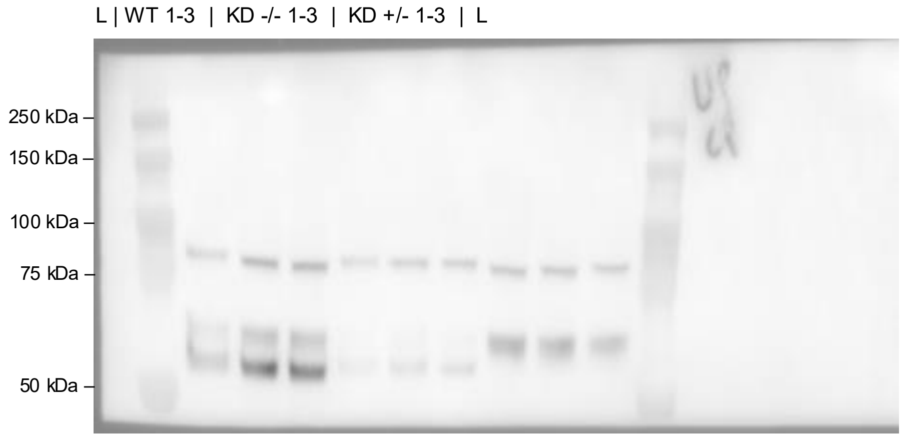


**CPSF7 KD & WT: GAPDH (same membrane)**

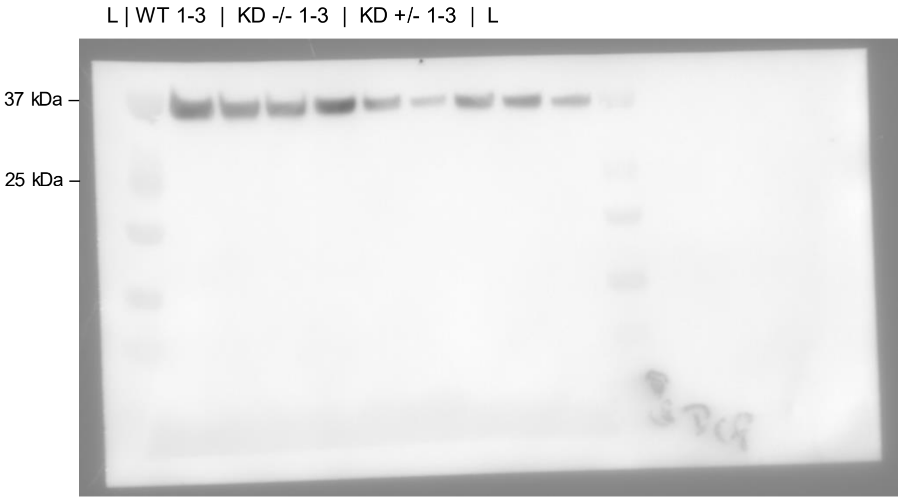


Uncropped western blots for Extended Data Figure 5I (continued).
Each membrane was probed for the indicated RBP and then for the loading control (GAPDH) on the same membrane.
Lanes: WT, homozygous (-/-) and heterozygous (+/-) KD, n = 3 independent experiments each.
L, protein ladder (10-250 kDa, Precision Plus Protein Dual Color Standards #1610374). The CPSF7 antibody detects the characteristic 52/56 kDa doublet.

### Supplementary Figure 1c: Uncropped gel for Extended Data Figure 7F

APP770-specific RT-PCR in RBP knockdown neurons

Row 1 - Experiment 1: L | CELF2 – SRSF5 – SUPT6H – HNRNPLL - SCAF11 – TIA1 - TIAL1 – CPSF7 – WT Experiment 2: L | same order
Row 2 - Experiment 3 with technical duplicates: L | CELF2 ×2 – SRSF5 ×2 – SUPT6H ×2 – HNRNPLL ×2 - SCAF11 ×2 – TIA1 ×2 - TIAL1 ×2 – CPSF7 ×2 – WT ×2 | L

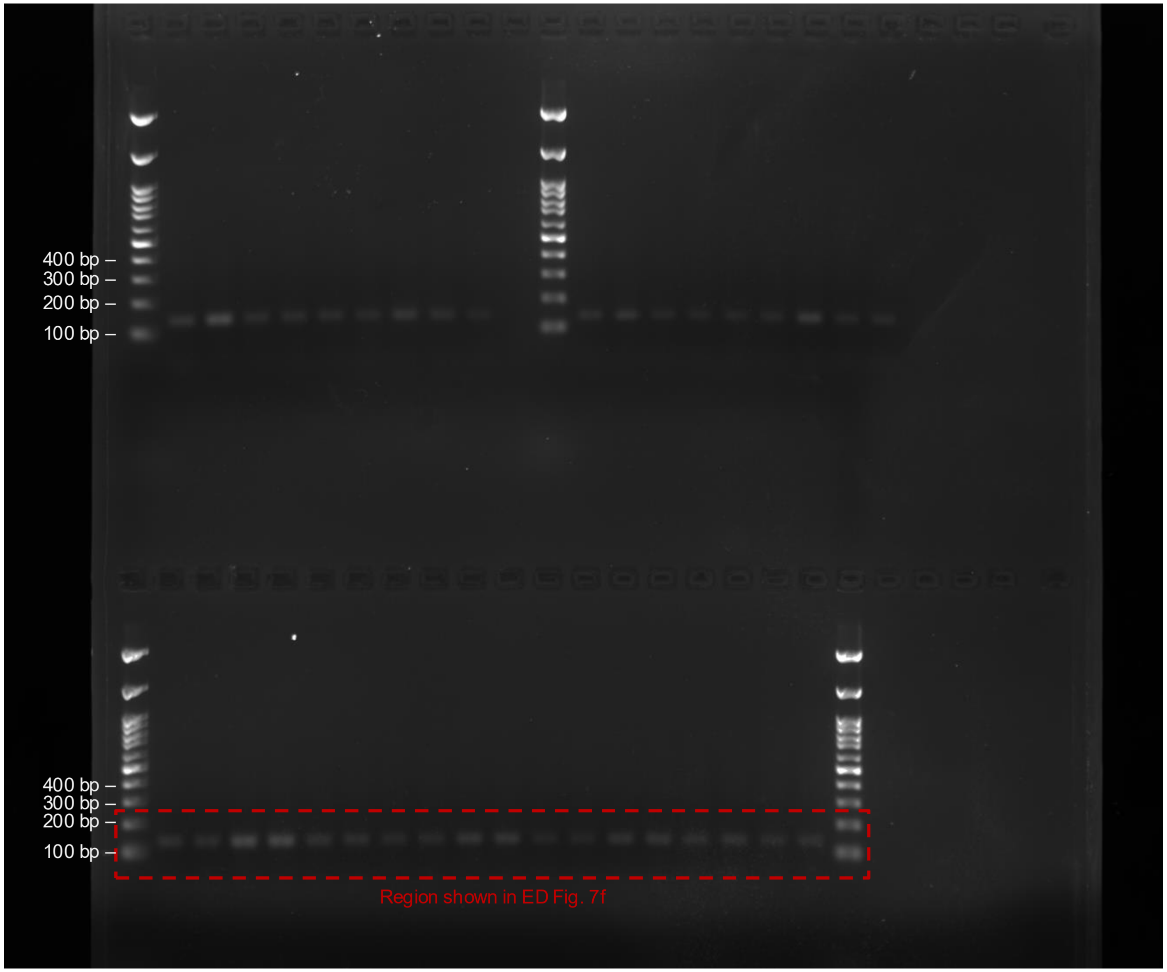


Uncropped image of the RT-PCR gel quantified in Extended Data Figure 7F (APP770-specific product, 136 bp).
Upper row: independent experiments 1 and 2. Lower row: independent experiment 3, loaded in technical duplicate, which was ave raged before quantification (n = 3 independent experiments).
Lanes were loaded in the order CELF2, SRSF5, SUPT6H, HNRNPLL, SCAF11, TIA1, TIAL1, CPSF7 and WT. The image shown in Extended Data Figure 7F was flipped horizontally so that WT appears first.
L, 100 bp ladder (first and middle wells, upper row; first and last wells, lower row). The dashed box indicates the region shown in Extended Data Figure 7F.
